# *Pinus* sp. leaf extracts exert antileishmanial effects against *Leishmania donovani* by targeting trypanothione reductase

**DOI:** 10.64898/2026.04.08.717320

**Authors:** Raoul Kemzeu, Lauve Rachel Tchokouaha Yamthe, Cyrille Armel Njanpa Ngansop, Eugenie Aimee Madiesse Kemgne, Boniface Pone Kamdem, Vincent Ngouana, Alvine Ngoutane Mfopa, Canis Parfait Donbou Djiotie, Patrick Valere Tsouh Fokou, Jaures Marius Tsakem Nangap, Lunga Paul Keilah, Lenta Ndjakou Bruno, Fabrice Fekam Boyom

## Abstract

The urgent need for new antileishmanial drugs with novel mechanisms of action is driven by the limited efficacy, high toxicity, and growing resistance to currently available treatments. This study investigates the antileishmanial potential of extracts from *Pinus* sp., a plant widely used in the Cameroonian pharmacopoeia to treat various infections, including cough and fever conditions. The antileishmanial activity of the ethanol, methanol, hydro-ethanol, and hydro-methanol extracts of *Pinus sp.* leaves were evaluated against *Leishmania donovani* promastigotes and amastigotes using a resazurin-based assay. The most active extract was then subjected to UHPLC-LC-MS/MS metabolomic profiling. Cytotoxicity, immunomodulatory, antioxidant, and anti-inflammatory properties of *Pinus sp.* extracts were also investigated. Moreover, the pharmacokinetic profile and molecular interactions of the identified compounds were predicted through *in silico* experiments using trypanothione reductase as the target enzyme. The acute toxicity study of the most promising extract (ethanol extract) was performed at 2000 and 5000 mg/kg using the organization for economic co-operation and development (OECD) guidelines, number 423. The plant extracts inhibited the growth of *L. donovani* with IC_50_ values ranging from 6.45 to 15.78 µg/mL and 16.11 to 24.44 µg/mL in promastigotes and amastigotes, respectively. These extracts also showed antioxidant, immunomodulatory, and anti-inflammatory effects. The extracts indicated no cytotoxicity and high selectivity (SI >10) toward Raw264.7 and Vero CRL1586 cells. The ethanolic extract showed a lethal dose (LD₅₀) value greater than 5000 mg/kg, thus highlighting its non toxicity upon the acute toxicity test. The metabolomic profiling of the ethanol extract identified several compound classes, including flavanone O-glycosides (e.g. epiafzelechin trimethyl ether), alkaloids (e.g. harmane), and diterpenes (e.g. abietic acid). Further molecular docking studies revealed strong binding affinities of these compounds to trypanothione reductase, thus suggesting the inhibitory potential of these compounds toward the target enzyme. Overall, extracts from *Pinus* sp. leaves demonstrate promising antileishmanial effects. However, isolation and characterization of the antileishmanial compounds from *Pinus sp.* leaf, detailed pharmacokinetics, and mechanisms of action are warranted for the successful utilization of this plant in antileishmanial drug discovery.

## 1. INTRODUCTION

Leishmaniases are a group of neglected tropical diseases caused by protozoan parasites of the genus *Leishmania*. These parasites are transmitted to humans through the bite of an infected female sandfly from the genus *Phlebotomus* (Abdoul-Latif et al., 2024; Djune Yemeli et al., 2021). Of the more than 30 *Leishmania* species known to infect mammals, approximately 20-21 species are recognized as pathogenic to humans, causing three main clinical forms of the disease, including cutaneous, mucocutaneous, and visceral leishmaniasis. The latter form is the most severe and is fatal in over 95% to 100% of cases if left untreated (Abdossamadi et al., 2017; Nourian et al., 2019). According to the World Health Organization, leishmaniasis is a serious public health problem in over 100 countries, with approximately 12 million people infected and over 350 million at risk (Abdoul-Latif et al., 2024). In Cameroon, both cutaneous and visceral leishmaniasis have been reported since 2001, particularly in the far north region (Mokolo) and the border areas with Chad (Ngouateu & Dondji, 2022; Tateng et al., 2024).

Leishmaniasis pathogenesis involves the injection of *Leishmania* promastigotes by sandflies, which are phagocytized by macrophages and transform into intracellular amastigotes (Gupta et al., 2022; Loeuillet et al., 2016). The promastigote form exists in the sand fly vector, where it undergoes various differentiation steps and transforms into the infective metacyclic promastigote form. These metacyclics are transmitted to mammalian hosts during the bite of the sand fly (Gupta et al., 2022). Amastigotes are the forms within the mammalian host, especially in phagocytic cells where they survive as intracellular parasites. When a sand fly vector bites an infected mammal, it ingests the amastigotes, which transform into the flagellated promastigote form on reaching the midgut of the insect. Eventually, the promastigotes move to the alimentary tract of the insect where they survive extracellularly and multiply by binary fission. The promastigotes then migrate towards the salivary glands and oesophagus and are later transmitted along with the insect saliva to the mammalian host during the next blood meal (Gupta et al., 2022). During infection, macrophages are the primary host cells, and their ability to control the parasite largely depends on the production of reactive oxygen species and nitric oxide (NO) (Horta et al., 2012; Roy et al., 2017). *Leishmania* parasites rely on trypanothione reductase (TR) to maintain redox balance and survive the harsh, oxidizing environment of host macrophage phagolysosomes. Trypanothione reductase is a well-validated, prime target for drug development against kinetoplastid parasites (e.g. *Leishmania* and *Trypanosoma*) because it is essential for their survival, protecting them from oxidative stress (Al-Khalaifah, 2022; Swenerton et al., 2010). Unlike humans, who rely on the glutathione reductase (GR) system, these parasites exclusively use the trypanothione system, making TR a target with high selectivity (Turcano et al., 2020). Mechanistically, inhibition of TR disrupts the unique redox metabolism of trypanosomatid parasites, leading to a significant increase in their sensitivity to oxidative stress, which ultimately results in programmed cell death or apoptosis (Fiorillo et al., 2022; Turcano et al., 2020). Accumulated evidence has shown a significant shift from “one-target-one-drug” to a more holistic approach in antileishmanial drug discovery given the rate of resistance to traditional drugs (Cemali et al., 2025). In fact, effective drug candidates for leishmaniasis are moving toward a polypharmacological approach, where a single compound simultaneously acts as an antiparasiticagent, inhibits specific essential enzymes, and modulates the host immune system to enhance Th1-type responses (Bora et al., 2024; Makarani et al., 2025). *In silico* methods, which involve using computer simulations, molecular modeling, and docking, are considered essential, cost-effective, and time-saving tools for accelerating the identification and optimization of new drug candidates (Cohen & Azas, 2021; Lamotte et al., 2019).

Current chemotherapy for leishmaniasis faces significant challenges, as the primary treatments pentavalent antimonials, amphotericin B, and miltefosine are hampered by toxicity, emerging drug resistance, high costs, and complex administration, often requiring hospitalization in resource-limited settings (Olias-Molero et al., 2021; Zhang et al., 2025). Thus, there is a pressing need to search for effective treatments against leishmaniasis.

Plants have served as a foundational source of medicine for millennia, and their complex, diverse, and often un-investigated chemical structures offer a rich, largely untapped reservoir for discovering novel active principles, with approximately 80% of people worldwide using traditional herbal medicines as a first line of defense. Examples of such plants include the *Pinus* species, which belong to the Pinaceae family, comprising over 100 species of evergreen conifers (Burley & Barnes, 2004). Traditional uses of *Pinus* species (pine trees) span medicine, food, and crafting, utilizing resin, needles, bark, and seeds for remedies against respiratory issues, fever conditions, and other infections (Burley & Barnes, 2004; Dziedzinski et al., 2021; Papp et al., 2022). Phytochemical screening of *Pinus* species reveals a rich profile of bioactive compounds, predominantly terpenes (α-pinene, β-pinene), phenolics, flavonoids, and tannins, with significant antioxidant, antibacterial, and antifungal activities (Piechowiak et al., 2023; Sharma et al., 2015).

Moreover, traditional uses of various *Pinus* species (pine) for treating fevers, colds, and associated infections have been documented across many cultures, often focusing on the use of needles, inner bark, and resin (Sharma et al., 2015). As a matter of fact, these plants are commonly employed in traditional medicine due to their diaphoretic (fever-reducing), antiseptic, and antioxidant properties, particularly in treating colds, coughs, and fever related symptoms (Papp et al., 2022). Thus, the scientific validation of *Pinus* species in treating fever related conditions, including leishmaniasis is of outstanding importance. Therefore, this study sought to evaluate the inhibitory effects of *Pinus* sp. extracts against promastigote and amastigote forms of *L. donovani*. The phytochemical analysis of the most promising extract, ADMET prediction of plant annotated compounds and their molecular docking toward trypanothione reductase are also investigated.

## 2. MATERIALS AND METHODS

### 2-1. Preparation of crude extracts and stock solutions

*Pinus* sp. leaves (Figure 1) were collected on 13th April, 2022, at the Campus 1 of the university of Yaounde 1, Ngoa-Ekelle, Yaounde III (3°51’26” Nord, 11°30’4”Est). The plant was identified at the National Herbarium of Cameroon where a voucher specimen was deposited under the reference number 50178/HNC. After plant collection, *Pinus sp.* leaves were dried at ambient temperature (29-32°C) and ground using a mechanical blinder. Five hundred grams of leaf powder were separately macerated at ambient temperature in 3 L of each solvent (ethanol, methanol, and water-ethanol and water-methanol mixtures (30:70, v/v) for 72 hours. The mixtures were gently shaken twice daily (morning and evening) to optimize extraction. Plant residues were macerated twice more consecutively with the same solvent volume. After 72 hours, each macerate was filtered using Whatman No. 1 filter paper and then evaporated using a rotary evaporator (BÜCHI 011) at 80°C for ethanol and hydroethanolic extracts, and at 70°C for methanol and hydromethanolic extracts. The extracts were dried under continuous fan ventilation in the laboratory for 5 days. The crude extracts obtained were weighed, and extraction yields were calculated as the percent ratio of extract mass to the starting plant material mass. Stock extract solutions were prepared at 100 mg/mL by dissolving 100 mg of each extract in 1 mL of 99% dimethylsulfoxide (DMSO), and then vortexed thoroughly in 1.5 mL Eppendorf tubes. So prepared extracts were stored at 4°C until further use for antileishmanial and cytotoxicity tests.

**Figure 1:**
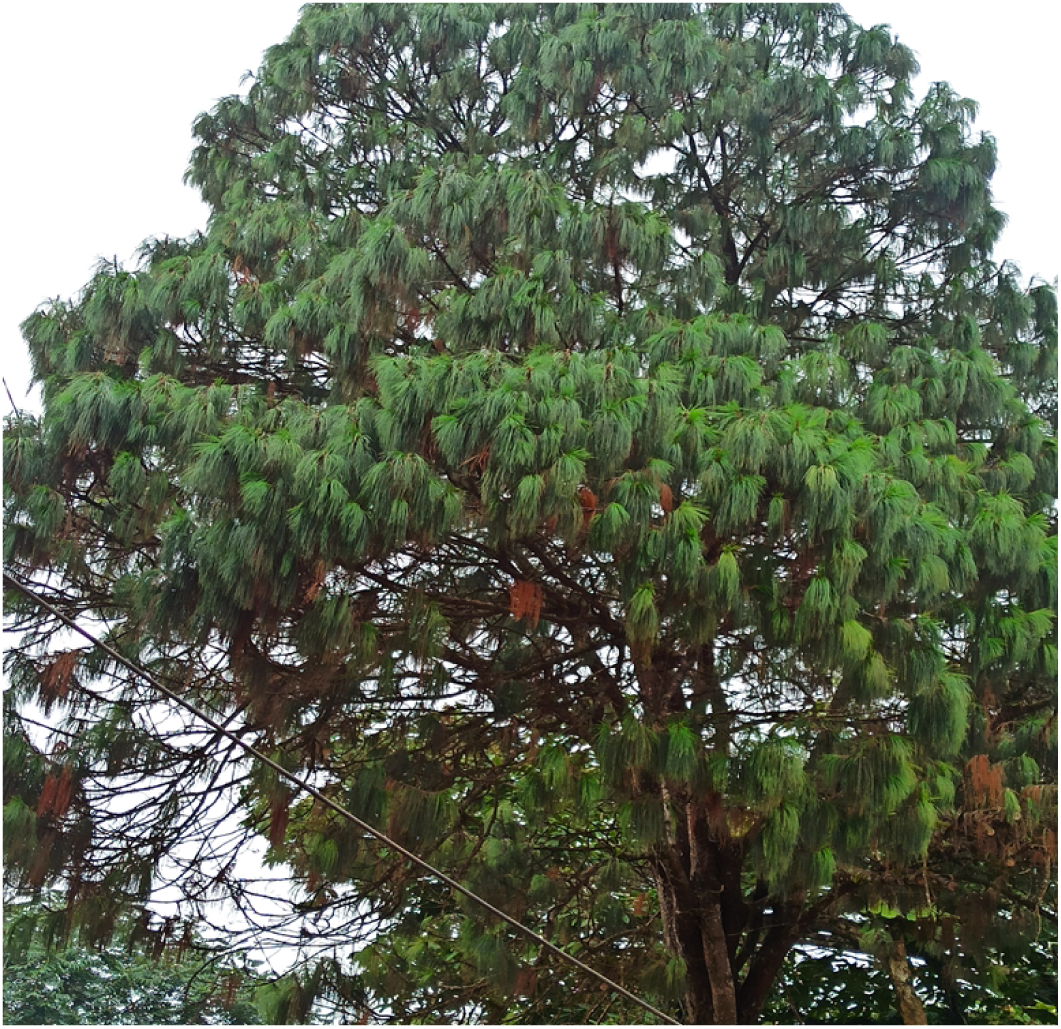
Photograpgh of Pinus sp. leaves (by R Kemzeu, 2022)

### 2-2. Antileishmanial and cytotoxicity tests

#### 2-2-1. Culture of *Leishmania donovani* and the normal mammalian cells

##### Leishmania donovani parasites culture

The cryopreserved promastigote form of *L. donovani* (1S (MHOM/SD/62/1S)) was obtained from BEI Resources (https://www.beiresources.org/) and routinely cultured at the Antimicrobial and Biocontrol Agents Unit (AmBcAU), University of Yaoundé I, in Medium 199 (Sigma, Darmstadt, Germany) supplemented with 10% heat-inactivated fetal bovine serum (HIFBS) (Sigma) and antibiotics (100 IU/mL penicillin and 100 μg/mL streptomycin). Cultures were maintained in 75 cm² flasks at 28°C, monitored daily for growth, and subcultured every 72 hours.

##### Mammalian cells culture

RAW 264.7 and VERO CRL1586 cells obtained from the American Type Culture Collection (ATCC) were cultured in 25 cm² flasks containing complete Dulbecco’s Modified Eagle’s Medium (DMEM) supplemented with 13.5 g/L DMEM powder (Sigma Aldrich), 10% fetal bovine serum (Sigma Aldrich), 0.2% sodium bicarbonate (w/v) (Sigma), and 50 μg/mL gentamicin (Sigma Aldrich). The culture medium was renewed every 72 hours. Upon reaching 70–80% confluence, cells were detached using 1 mL of 0.05% trypsin-EDTA solution for 5 minutes.

#### 2-2-2. *In vitro* assay for *L. donovani* promastigote inhibition

The antileishmanial activity was evaluated on *L. donovani* promastigotes using the resazurin colorimetric assay as described by (Siqueira-Neto et al., 2010). Briefly, 4×10⁵ promastigotes/mL/well were seeded in 96-well microtiter plates and treated with serially diluted concentration of the extracts at 100 to 0.16 µg/mL, at 28°C for 72 hours. Upon incubation, 1 mg/mL of resazurin (Sigma) was added in each plate well, and plates were incubated for an additional 44 hours at 28°C. Cell viability was determined by measuring the fluorescence proportional to the amount of pink resorufin produced through the reduction of blue resazurin by mitochondrial dehydrogenases of live parasites using a Tecan Infinite M200 multiwell plate reader (Tecan) with excitation/emission wavelengths set at 530/590 nm. Negative and positive controls consisted of 100 µL of parasites only and amphotericin B (Sigma) at 10 to 0.016 µg/mL, respectively. Half-maximal inhibitory concentration (IC₅₀) values were calculated using GraphPad Prism 8.0.1 (244) software, with dose-response curves fitted by non-linear regression.

#### 2-2-3. Cytotoxicity assay

Cytotoxicity was evaluated on RAW 264.7 and Vero CRL1586 cells using the Alamar blue assay as described by Bowling et al. (2012). Macrophages/Vero were seeded into 96-well flat-bottom plates at a density of 10⁴ cells/well in 100 μL of complete medium and incubated for 24 hours at 37°C, 5% CO₂, and 70% relative humidity to allow adhesion. Next, ten microliters of serially diluted test samples were added in triplicate, with concentrations ranging from 500 to 0.8 µg/mL, and plates were incubated for 48 hours under the same conditions. Growth control consisted of 0.1% DMSO (representing 100% cell growth), while the positive control podophyllotoxin (Sigma) was used at concentrations ranging from 20 to 0.032 μM. Cell proliferation was assessed by adding 10 μL of 0.15 mg/mL resazurin stock solution in sterile PBS to each well, followed by incubation for 4 hours. Fluorescence was measured on a Tecan Infinite M200 reader at excitation/emission wavelengths of 530/590 nm. Results were expressed as half-maximal cytotoxic concentration (CC_50_) values derived from dose-response curves using GraphPad Prism 8.0.1 (244) software. Selectivity indices (SI: CC_50_/IC_50_) were calculated to evaluate the therapeutic window.

#### 2-2-4. Inhibition of intracellular amastigote form of *L. donovani*

The activity of plant extracts against intracellular amastigotes was evaluated following Jain et al. (2012) with slight modifications (Jain et al., 2012). Briefly, RAW 264.7 macrophages (4×10³ cells/well) were seeded in 96-well plates and incubated for 6 h at 37°C, 5% CO₂ for adhesion. Non-adherent cells were removed by washing with sterile PBS. Adherent macrophages were infected with metacyclic promastigotes (4×10⁵ cells) at a 1:10 macrophage-to-parasite ratio and incubated for 24 h at 37°C, 5% CO₂ to allow infection. Subsequently, the medium was removed, and monolayers washed four times with PBS to eliminate free parasites. Fresh M199 medium supplemented with 10% FBS and test extracts at serially diluted concentrations (100 to 0.16 µg/mL) were added in triplicate wells and incubated for 48 h at 37°C, 5% CO₂. Following incubation, 0.05% sodium dodecyl sulfate (SDS) was added for 30 seconds to lyse macrophages in a controlled manner, followed by addition of M199 with 10% FBS to stop lysis. The negative and positive controls consisted respectively of cells treated with 100 µL of parasites without inhibitor and with amphotericin B (Sigma) at concentrations ranging from 10 to 0.016 µg/mL. Resazurin reagent (250 μg/mL) was added, and plates incubated for additional 24 h before fluorescence measurement (excitation, 530 nm; emission, 590 nm) using a Tecan Infinite M200 reader. Inhibition percentages were calculated using Microsoft Excel, and IC_50_ values derived from dose-response curves using GraphPad Prism 8.0.1 (244) software. The promastigote selectivity index (SPI = IC_50_ on promastigote form / IC_50_ on amastigote form) was also calculated to evaluate the relative efficacy against the two life cycle stages. Interpretation criteria as defined by (De Muylder et al., 2011) were: SPI > 2 indicates higher activity on amastigotes; SPI < 0.4 indicates higher activity on promastigotes; SPI between 0.4 and 2 indicates activity on both forms.

#### 2-2-5. Immunomodulatory and anti-inflammatory tests

##### 2-2-5-1. Immunomodulatory activity

The levels of nitric oxide (NO) produced by infected macrophages following treatment with test samples were determined using the Griess reaction (Sigma-Aldrich, USA), as previously described by Karampetsou et al. (2019). Following the anti-amastigote assay, 100 µL of supernatant were added to the 48 hour-treated cells in the 96-well plate. Next, 100 µL of a freshly prepared Griess reagent were introduced into the plate wells (Sun et al., 2003). After that, the plates were incubated for 15 minutes at room temperature, and then the absorbance of the preparation was measured using a Magellan Infinite M200 multi-well plate reader (Tecan) at 570 nm. Sodium nitrite (Sigma Aldrich) at 75 µg/mL prepared in distilled water served as the positive control, with final concentrations ranging from 0.29 to 75 µg/mL. A standard curve was constructed using optical densities versus concentrations to quantify NO produced upon treatment with samples. Results were expressed as molar concentrations calculated from the linear equation y = 0.0036x + 0.0788 with R² = 0.9962, obtained from the calibration curve plotted using Microsoft Excel 2016.

##### 2-2-5-2. Anti-inflammatory assay

The *in vitro* anti-inflammatory activity was assessed by the bovine serum albumin (BSA) protein denaturation method, as described by Chandra et al. (Chandra et al., 2012). Briefly, 10 µL of BSA solution (40 mg/mL in phosphate buffer) was added to 140 µL of PBS (pH 7.4) in a 96-well microplate. Afterward, 100 µL of extract or diclofenac sodium at various concentrations (final volume = 2.5 µL) were added to the microplate. Methanol (100 µL) was used as a blank. Plant extracts and diclofenac sodium were tested at final concentrations of 100, 50, 25, 12.5, 6.25, 3.125 and 1.5625 µg/mL. After mixing the preparation, plates were incubated at 37°C for 30 minutes, then heated in a water bath at 70°C for 15 minutes. After cooling the preparations, the optical densities (OD) were read at 660 nm using a Tecan M200 multi-well plate reader. Tests were performed in duplicate. The percentage inhibition of protein denaturation was calculated using Microsoft Excel 2016, and IC_50_ values were determined with GraphPad Prism 8.0.1 (244) software.

### 2-2-6. Phytochemical screening of plant extracts

#### 2-2-6-1. Total phenolic content determination

Total phenolic content was determined spectrophotometrically as described by Singleton et al. (Singleton et al., 1999). Briefly, 200 µL of sample solution (100 mg/mL) and 100 µL of diluted Folin-Ciocalteu reagent (1:10) were added to test tubes. Then the tubes were shaken for 5 minutes, followed by an addition of 800 µL of sodium carbonate solution (7.5% Na₂CO₃). After that, the preparation was incubated in the dark for 1 hour, and the absorbance was measured against the blank at 765 nm. The quantification of the polyphenols was done based on a linear calibration curve (y=2.6156x-0.0153; r2 =0.9997) produced by the standard (gallic acid) at different concentrations (1000, 500, 250, 125, 62.5 and 31.25 µg/mL). The results were expressed in milligrams of gallic acid equivalent (GAE) per gram of dry extract (μg GAE/g DE) using the following formula: P = (C×V)/M, where P = phenolic content (mg GAE/g of dry extract), C = concentration of phenolic content obtained from calibration curve (mg/mL), V = volume (mL), and M = weight of dry extract (g).

#### 2-2-6-2. Total flavonoid content determination

The flavonoid content was determined as described by Marinova et al. (2005). Briefly, 400 µL of extract sample (250 µg/mL) was mixed to 120 µL of 5% NaNO₂ and allowed to stand for 5 minutes. Subsequently, 120 μL of 10% aluminium chloride was added, and the preparation was incubated for 6 minutes at room temperature. Then, 800 μL of 1 M sodium hydroxide was added, and the final volume was adjusted to 2.5 mL with distilled water. The absorbance of the preparation was measured at 510 nm using a Tecan M200 multi-well plate reader. A calibration curve (y = 0.2019x + 0.0012; r2 =0.9998) was drawn using catechin standards (1000, 500, 250, 125, 62.5 and 31.25 µg/mL) under the same conditions. The flavonoids content was calculated from the calibration curve and expressed in milligram of quercetin equivalent per gram of dry weight (mg QE/g DE) using the following formula:

P = (C×V)/M, where P = flavonoid content (mg GAE/g dry extract), C = concentration of flavonoid content obtained from the calibration curve (mg/mL), V = volume (mL), and M = weight of dry extract (g).

### 2-2-7. Antioxidant assay

#### 2-2-7-1. DPPH radical scavenging test

The free radical scavenging ability of the promising extracts was evaluated using the DPPH radical (2,2-diphenyl-1-picrylhydrazyl) as described by Bassene (Bassene, 2012). In brief, 75 µL of a DPPH solution (0.02%; w/v), prepared in methanol was added to 25 µL of different concentrations of extract (500, 250, 125, 62.5, 31.25, 15.62 and 7.81 μg/mL), followed by an incubation of the preparation in the dark for 30 minutes. After incubation, the absorbance was measured at 517 nm using a Tecan M200 multi-well plate reader. Negative and positive controls consisted of DPPH (100 µL) solution only and ascorbic acid (25, 12.5, 6.25, 3.125, 1.56, 0.78 and 0.39 μg/mL), respectively. The scavenging activity was calculated as the percentage of DPPH radical inhibition using the following formula: % Inhibition = [(Ac − Ae)/Ac]×100, where Ac is the absorbance of the negative control and Ae is the absorbance of the test extract. The IC_50,_ defined as the concentration of extract required to scavenge 50% of DPPH free radicals, was determined using GraphPad Prism 8.0.1 (244) software.

#### 2-2-7-2. ABTS radical scavenging assay

The ABTS (2,2’-azino-bis(3-ethylbenzothiazoline-6-sulfonic acid)) radical scavenging activity was assessed as per Khan et al.’s (Khan et al., 2012) protocol with minor modifications. The ABTS solution was prepared by mixing equal volumes of 7 mM ABTS solution in methanol and 4.9 mM potassium persulfate in distilled water. After that, the preparation was incubated in the dark at room temperature for 15 hours to produce a dark-colored solution. Prior to the test, the solution was diluted 20-fold with distilled water to obtain a final concentration of 0.35 mM. Then, 25 µL of each sample solution (500, 250, 125, 62.5, 31.25, 15.62 and 7.81 μg/ mL) was added in triplicate to 75 µL of 0.175 mM of the ABTS+ solution and incubated for 30 minutes at room temperature in the dark. The decrease in the absorbance of ABTS was measured at 734 nm using a Tecan M200 multi-well plate reader. Ethanol was used as a blank, whereas the ABTS solution served as the negative control. Ascorbic acid (25, 12.5, 6.25, 3.125, 1.56, 0.78 and 0.39 μg/mL) was used as the positive control. The percentage of inhibition was calculated as follows: % Inhibition = [(Ac−Ae)/Ac]×100, where Ac and Ae are the absorbances of negative control and test sample, respectively. The IC_50_ values were determined using the GraphPad Prism 8.0.1 (244) software.

#### 2-2-7-3. Ferric reducing antioxidant power test

This test is based on the ability of extracts to reduce Fe³⁺ ions to Fe²⁺ ions, which form a red-orange complex with 1,10-phenanthroline, measurable at 505 nm. The ferric reducing power was evaluated following the protocol described by Canada (Canada, 1994). Equal volumes (25 µL) of extracts (2000 µg/mL) and the Fe³⁺ solution (1.2 mg/mL in distilled water) in the 96 well plates. After that, plates were incubated for 15 minutes at room temperature, then 50 µL of 0.2% orthophenanthroline in methanol was added, followed by an additional incubation for 15 minutes. Optical densities were read at 505 nm using the Tecan M200 multi-well plate reader. Negative control (0% reduction) consisted of 25 µL of methanol, 25 µL Fe³⁺ solution, and 50 µL of orthophenanthroline, the positive control (100% reduction) consisted of 25 µL of Fe³⁺ solution, 25 µL ascorbic acid and 50 µL of orthophenanthroline solution. The test concentrations were (500, 250, 125, 62.5, 31.25, 15.62 and 7.81 μg/ mL) and (25, 12.5, 6.25, 3.125, 1.56, 0.78 and 0.39 μg/mL) for extract and positive control, respectively. The reducing capacity for the Fe³⁺ solution is directly proportional to the optical density. The percentage of reduction of each sample is calculated based on the activity of ascorbic acid (100% reduction of Fe³⁺ solution). The IC_50_ values were further calculated using GraphPad Prism 8.0.1 (244) software.

### 2-2-7. *In vivo* acute oral toxicity of *Pinus sp.* ethanol extract

The acute oral toxicity was carried out using female albino Wistar rats aged from 10 to 12 weeks, weighing from 180 to 220 g. The test was performed according to the OECD guideline 423 (OECD, 2001). Approval for animal studies was obtained from the Institutional Ethical Committee (Cameroon), in compliance with all procedures recommended by the European Union on the protection of animals used for scientific purposes (CEE Council 86/609; Ref N° FWA-IRD 0001954). The albino Wistar rats were bred in the animal house of the Laboratory for Phytobiochemistry and Medicinal Plants Study of the University of Yaounde I (Cameroon). The animals were kept at room temperature with a natural nictemeral cycle. They were divided into three groups of three animals each. Groups 1 and 2 received a single oral dose of the extract (1 mL/100 g body weight) at 2000 mg/kg and 5000 mg/kg, respectively. Group 3 served as a negative control and the animals of this group received the vehicle control (10% Tween 80). After 0.5 and 4 hours post-treatment, animals were monitored for clinical signs of toxicity, body weight changes and mortality. This was followed by a 14 days’ observation to identify additional signs and symptoms of toxicity. After the observation time period had elapsed animals were fasted for 24 hours without water, then sacrificed by cervical dislocation to collect different organs (liver, kidneys, heart, lungs and spleen) that further helped to carry out biochemical and histological analyses.

Changes in the percentage of body weight and relative organ weights were calculated as follows:

Changes in the percentage of body weight (P) = ((Pi − P₀)/P₀)×100.

Relative organ weight (W) = (Po/Pr)×100, where P₀ = Body weight on day 0, Pi = Body weight 24-hour prior to test (fasting day), Po = Organ weight, and Pr = Body weight of rats on the fasting day prior to sacrifice.

Hematological parameters, including red blood cells (RBC), hematocrit (HCT), hemoglobin (HGB), white blood cells (WBC), platelets (PLT), mean corpuscular volume (MCV), lymphocytes (LYM), monocytes (MON), and granulocytes (GRA) were measured using an automated analyzer (Mindray BC-5300, Italia). Hepatic toxicity markers (alanine aminotransferase (ALT), aspartate aminotransferase (AST), alkaline phosphatase (ALP) and renal toxicity markers (total proteins, creatinine, uric acid) were assessed using respective kits (LABKIT, Spain) (Jacques & Brice, 2020; Smith & Bruton, 1977; Young, 2001). For histological analyses, liver and kidneys of test animals, which were introduced into 10% buffered formalin, were dehydrated by successive passages in graded concentrations of alcohol and then embedded in paraffin. A series of 5 μm paraffin sections were stained with haematoxylin and eosin (HE) for examination under an Olympus light microscope and photography at 20X ocular and 10X objective (HE × 200). The statistical analysis was carried out using analysis of variance (ANOVA), followed by Tukey’s post-test for the separation of means with GraphPad Prism 8.0.1 (244) software.

### 2-2-8. UHPLC-MS analysis of the promising extract

High-resolution mass spectra were acquired using a QTOF (quadrupole time-of-flight) mass spectrometer (Bruker, Germany) equipped with a HESI (heated electrospray ionization) source operating in negative ion mode. The instrument was configured to scan within a mass range of 100–1500 m/z at a rate of 1.00 Hz, ensuring high-accuracy mass measurements with a deviation of 0.40 ppm using sodium formate as a calibrant. The experimental parameters included a spray voltage of 4.5 kV, a capillary temperature of 200 ^◦^C, and a nitrogen sheath gas flow rate of 10 L/min. The QTOF mass spectrometer was coupled to an Ultimate 3000 UHPLC system (Thermo Fisher, USA) consisting of an LC pump, a Diode Array Detector (DAD) operating in the wavelength range of 190–600 nm, an autosampler with a 10 μL injection volume, and a column oven set at 35^◦^C. Chromatographic separation was performed on a Synergy MAX-RP 100A column (50×2 mm, 2.5 μ size particle) using water (+0.1 % formic acid) (solvent A)/acetonitrile (+0.1 % formic acid) (solvent B) gradient at a flow rate of 500 μL/min and an injection volume of 5 μL. The gradient program involved an initial isocratic elution with 95% A for 1.5 min, followed by a linear gradient to 100% B over 6 min, an isocratic elution with 100% B for 2 min, a 1 min return to the initial conditions (90% A), and a final equilibration period lasting for a minute. The acquired mass spectra were converted to the mzML format before being processed using MZmine 3.9 software. This allowed for visualization of the total ion chromatogram (TIC) and further conversion to the.abf format for subsequent analysis. The metabolites were then annotated based on their accurate mass, fragmentation patterns, and retention time by searching against several databases, including MASSBANK, MetaboBASE, BioSilico, PFAS-TOX, and the Exploit Database (PUBLIC-EXP). To facilitate this process, an in-house database in.msp format was uploaded into the MS-DIAL ware version 5 to annotate the metabolites.

### 2-2-9. *In Silico* studies

#### 2-2-9-1. Prediction of ADME and toxicity parameters of annotated compounds

The artificial intelligence-driven platform ADMET-AI (https://admet.ai) was used to predict the ADME (Absorption, Distribution, Metabolism, and Excretion) and toxicity profiles of the phytochemical constituents of the most promising extract. This platform utilizes advanced machine learning models trained on large ADMET datasets to rapidly screen and evaluate potential hit compounds. The structures of the identified metabolites were initially drawn using ChemBio2D Draw software. The corresponding SMILES (Simplified Molecular Input Line Entry System) codes for each compound were then generated and used as an input for the ADMET-AI web interface. The platform was used to predict key ADME parameters, including oral drug absorption, human intestinal absorption, skin permeability, and transdermal absorption.

The predictive power of ADMET-AI is based on its graph neural network architecture, known as Chemprop-RDKit. These models were trained on 41 ADMET datasets from the Therapeutics Data Commons (TDC) and have demonstrated top average rankings in the TDC ADMET Benchmark Group. This robust methodology ensures that ADMET-AI provides a scientifically sound and efficient approach for evaluating the ADMET profile of potential hit compounds.

#### 2-2-9-2. Molecular Docking

The molecular docking of the promising compounds was performed on *Leishmania infantum* trypanothione reductase as the target enzyme. Amphotericin B was used as the reference drug. This experiment was done as per a previously described procedure (Ayodele et al., 2023; Ibrahim et al., 2020; Kannan et al., 2024). In brief, the assay was performed in three main steps.

Firstly, the crystal structure of the target enzyme, *L. infantum* trypanothione reductase complexed with an inhibitor (PDB ID: 6ER5), was downloaded from the Protein Data Bank (https://www.rcsb.org/). This structure was selected because the presence of a co-crystallized ligand provided a well-defined active site. The target enzyme was prepared using PyRx Virtual Screening Tools, which involved removing the co-crystallized ligand and all water molecules, adding polar hydrogens, and converting the protein structure into the required AutoDock Vina pdbqt format. The binding site for docking was defined by a grid box centered on the original co-crystallized ligand.

Next, the 3D chemical structures of the potential hit compounds, which were identified by UHPLC-LC-MS/MS, were drawn using ChemDraw Professional 15.0 and Chem3D 15.0. These compound structures were subjected to PyRx for energy minimization using the MMFF94 force field and for converting into Autodock pdbqt ligand format.

Lastly, molecular docking simulations were conducted using AutoDock Vina, integrated within the PyRx software. The molecular docking was used to predict the binding affinity (in Kcal/moL) and the most favorable pose of each ligand within the enzyme’s active site. Then, the resulting docking poses and molecular interactions were analyzed and visualized using Discovery Studio 2021 software. This analysis allowed for the identification of key amino acid residues and specific chemical bonds (e.g., hydrogen bonds, hydrophobic interactions, and π-stacking) involved in the ligand-protein interactions.

## 3. RESULTS AND DISCUSSION

### 3-1. RESULTS

#### 3-1-1. Inhibitory effects of extracts against *L. donovani* and parasite selectivity

The inhibitory effects of the *Pinus sp.* extracts were evaluated against both the promastigote and intracellular amastigote forms of *L. donovani*, and their selectivity was assessed on Vero and RAW 264.7 cell lines. *Pinus sp.* extracts showed dose-dependent inhibition of the parasite’s viability, as illustrated by the dose-response curves (Figure 2). The incubation of amastigotes and promastigotes of *L. donovani* with different extracts of *Pinus sp.* led to a concentration-dependent parasite inhibition (Figure 2A). Moreover, there was a high percentage of growth when the normal cells Raw and Vero were incubated with extracts and podophyllotoxin (Figure 2B, Table 1).

**Figure 2:**
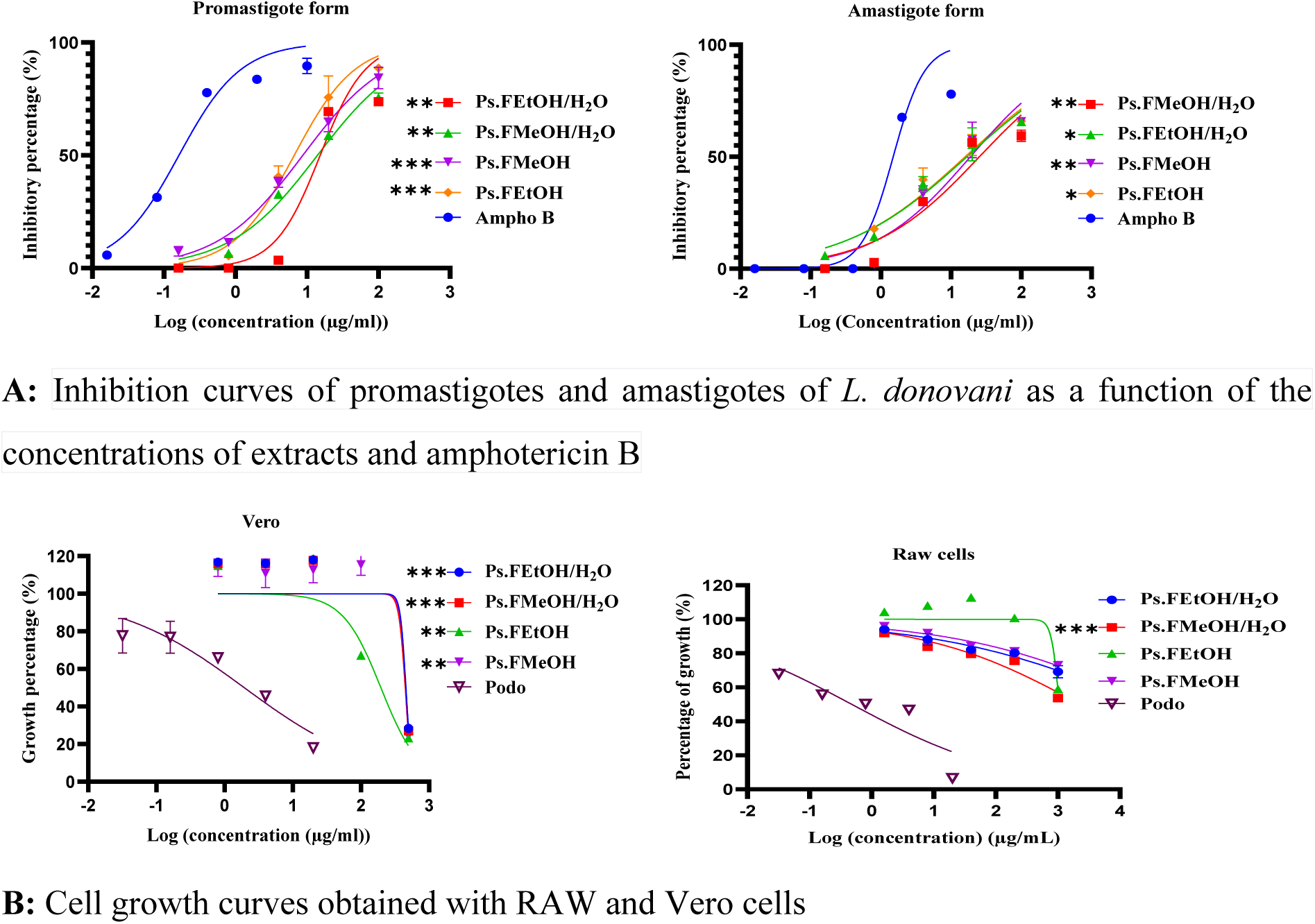
Parasite growth inhibition (A) and normal cell growth (B) curves *P ≤ 0.05, **P ≤ 0.01, ***P ≤ 0.001 (Dunnett’s test); Values are significantly different, compared with the value of the positive controls amphotericin B and podophyllotoxin; Ampho B: amphotericin B; Podo: podophyllotoxin.

**Table 1:** IC_50_ values of *Pinus sp*. extracts upon incubation with *L. donovani*, selectivity and specificity indices *P ≤ 0.05, **P ≤ 0.01, ***P ≤ 0.001, ****P ≤ 0.0001 (Dunnett’s test); Values are significantly different, compared to the value of the positive control amphotericin B; Ps.FEtOH: Ethanol extract of *Pinus sp*. leaves; Ps.FMeOH: Methanol extract of *Pinus sp*. leaves; Ps.FEtOH/H_2_O: Hydro-ethanolic extract of *Pinus sp*. leaves (30:70, v/v); Ps.FMeOH/H_2_O: Hydro-methanolic extract of *Pinus sp*. leaves (30:70, v/v); CC_50_ : 50% cytotoxic concentration ; IC_50_: Concentration that inhibits 50% of parasite growth; SD: Standard deviation; PC: Positive control; ND: Not determined; SI: Selectivity index; SPI: Specificity index.

| Extracts / Compounds | Yield of extraction (%) | CC<sub>50</sub> (µg/mL) |  | IC<sub>50</sub>±SD (µg/mL) and selectivity index |  |  |  |  |  | SPI |
| --- | --- | --- | --- | --- | --- | --- | --- | --- | --- | --- |
|  |  | Vero cells | Raw cells | Promastigotes | SI (Vero) | SI (Raw) | Amastigotes | SI (Vero) | SI (Raw) |  |
| **Ps.FEtOH** | 21.6 | 156.7±2.19 | >1000 | 9.67±2.28*** | 16.20* | >103.41 | 16.11±1.20* | 9.72** | >62.07 | 0.60±1.9 |
| **Ps.FMeOH** | 20.8 | 255.4±2.40 | >1000 | 6.45±0.002*** | 39.59* | >155.03 | 18.83±1.27** | 13.56* | >53.10 | 0.34±0.001 |
| **Ps.FEtOH/H<sub>2</sub>O (70/30)** | 9.4 | 235.2±2.37 | >1000 | 15.36±0.88** | 15.31* | >65.10 | 16.02±1.20* | 14.68** | >62.42 | 0.95±0.73 |
| **Ps.FMeOH/H<sub>2</sub>O (70/30)** | 11.6 | 228.5±2.35 | 922.25 | 15.78±0.58** | 14.48** | 58.44*** | 24.44±1.38** | 9.34** | 37.73* | 0.64±0.42 |
| **Amphotericin B** | ND | 36.68±16 | >200 | 0.15±0.002 | 244.53 | >1333 | 1.50±0.17 | 24.45 | >133.33 | 0.1±0.01 |
| **Podophyllotoxin** | ND | 1.951±0.29 | 1.69±0.31 | ND | ND | ND | ND | ND | ND | ND |

At 100 µg/mL, ethanol (Ps.FEtOH), methanol (Ps.FMeOH), hydroethanolic (Ps.FEtOH/H_2_O) and hydromethanolic (Ps.FMeOH/H_2_O) extracts from *Pinus sp*. showed significant (p<0.05) inhibition of *L. donovani* promastigotes with percentages ranging from 73.78% (for Ps.FEtOH/H_2_O) to 88.74% (for Ps.FMeOH).. The methanol extract (Ps.FMeOH) revealed the highest activity (IC_50_ : 6.45±0.002 µg/mL) when tested against *L. donovani* promastigotes. Against the intracellular amastigotes, IC_50_ values of 16.11±1.20, 18.83±1.27, 16.02±1.20 and 24.44±1.38 µg/mL were recorded for Ps.FEtOH, Ps.MeOH, Ps.FEtOH/H_2_O, and Ps.FMeOH/H_2_O extracts, respectively. The promastigote form was more susceptible to the methanol extract (Ps.FMeOH; SPI = 0.34), however; Ps.FEtOH/H_2_O (SPI = 0.66) and Ps.FEtOH (SPI = 1.52) revealed comparable activity on both forms of *L. donovani*.

Irrespective of the extracts tested and the cell lines used, the selectivity index was greater than 10, thus highlighting a high therapeutic window of these extracts.

#### 3-1-2. Immunomodulatory effects

In the nitric oxide assays, methanol, hydroethanolic and hydromethanolic extracts increased the NO production in infected macrophages compared to untreated controls, thus indicating the immunomodulatory effects of these extracts. By contrast, the ethanol extract of *Pinus sp.* (Ps.FEtOH) did not stimulate the NO production (Figure 3).

**Figure 3:**
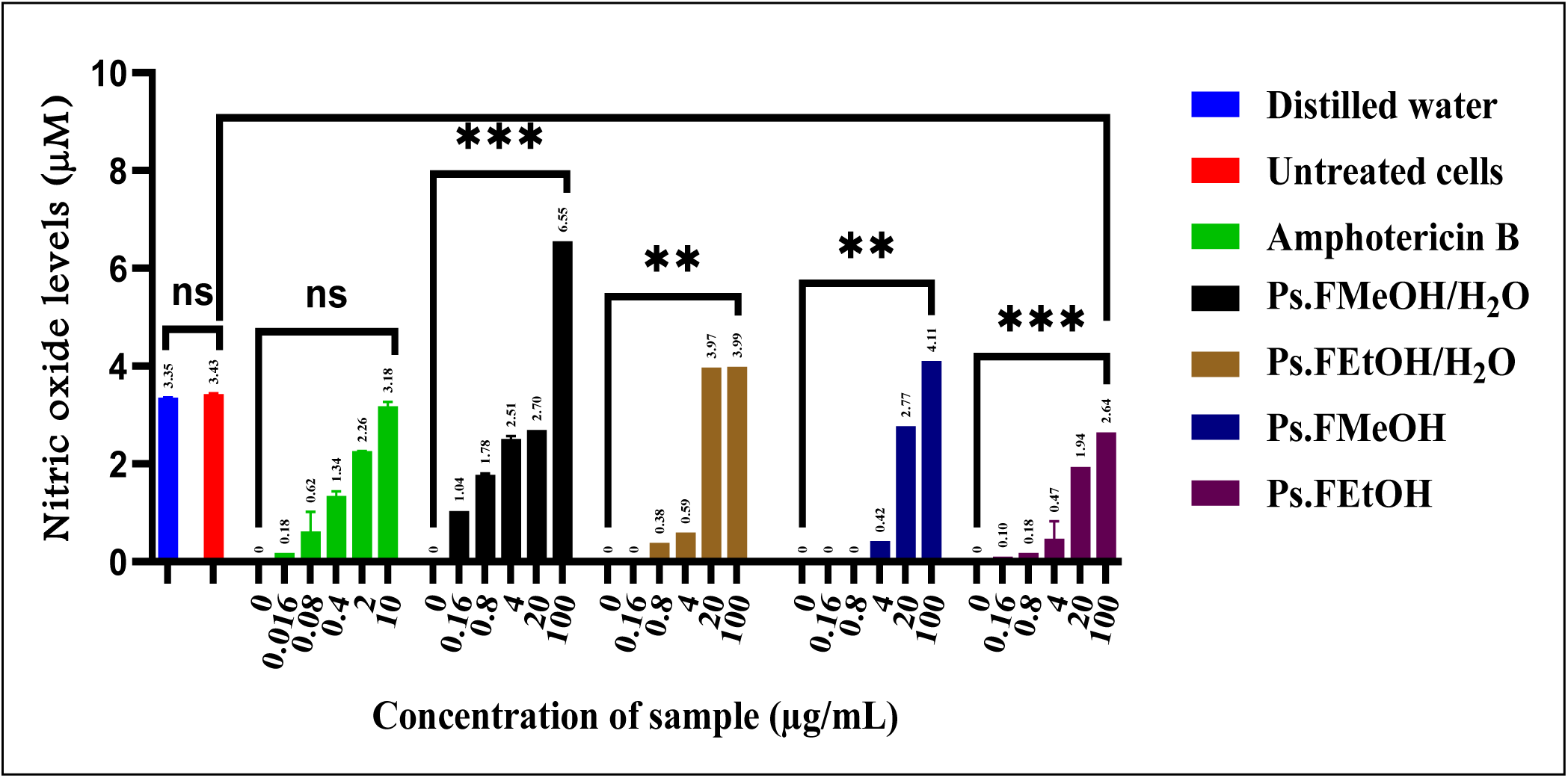
Quantification of nitric oxide upon macrophage cell treatment with different concentrations of *Pinus sp*. extracts and amphotericin B. **P ≤ 0.01, ***P ≤ 0.001, ns: Not significant (Dunnett’s test); Values are significantly different, compared to the value obtained for the positive control amphotericin B; Ps.FEtOH: Ethanol extract of *Pinus sp*. leaves; Ps.FMeOH: Methanol extract of *Pinus sp*. leaves; Ps.FEtOH/H_2_O: Hydro-ethanolic extract of *Pinus sp*. leaves (30:70, v/v); Ps.FMeOH/H_2_O: Hydro-methanolic extract of *Pinus sp*. leaves (30:70, v/v); µM: micromolar.

**Figure 4:**
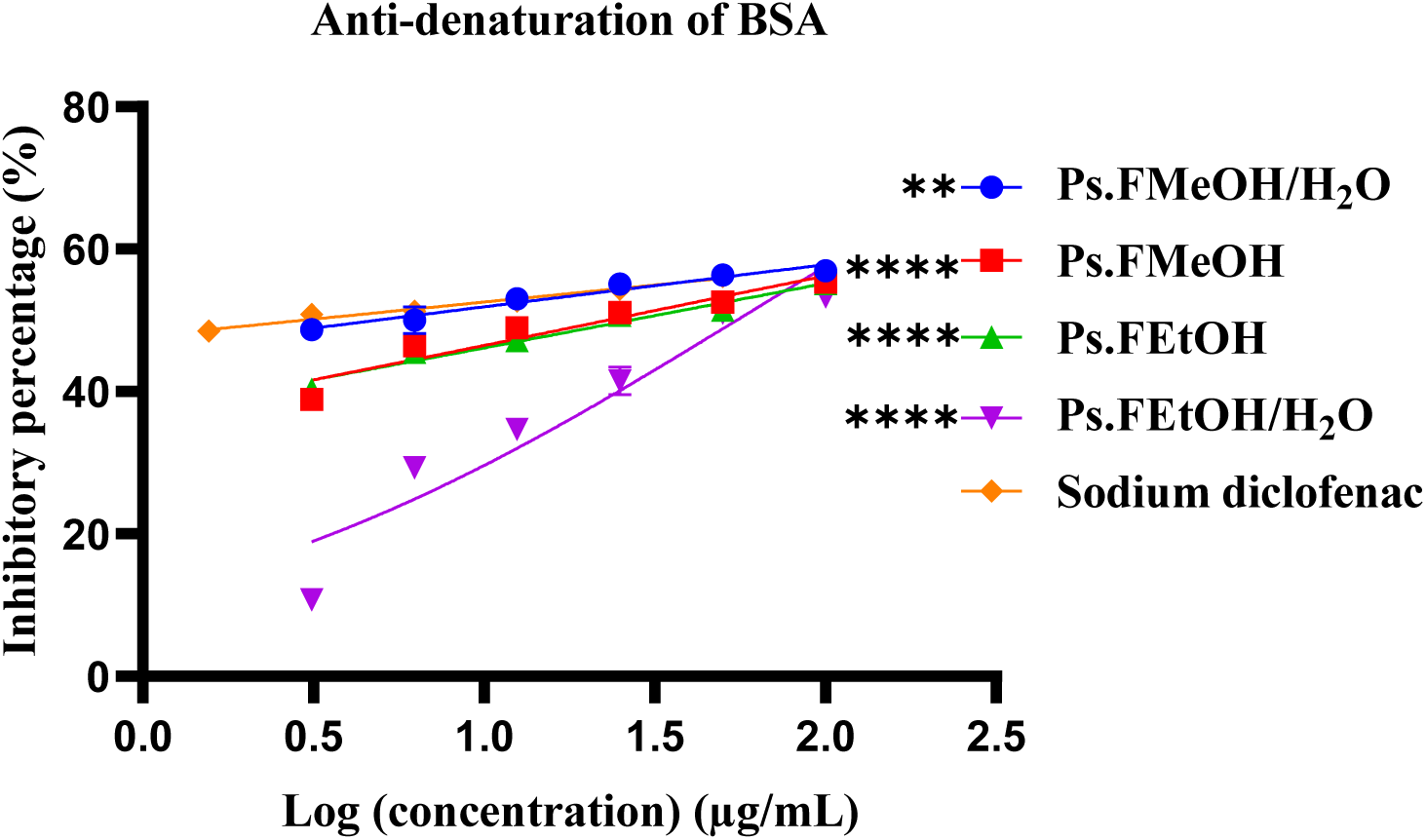
Percentages of inhibition of the BSA protein denaturation upon treatment with the *Pinus sp.* extracts and sodium diclofenac. **P ≤ 0.01, ****(P˂0.0001) (Dunnett’s test); Values are significantly different, compared to the value obtained for diclofenac sodium; Ps.FEtOH: Ethanol extract of *Pinus sp*. leaves; Ps.FMeOH: Methanol extract of *Pinus sp.* leaves; Ps.FEtOH/H_2_O: Hydro-ethanolic extract of *Pinus sp.* leaves (30:70, v/v); Ps.FMeOH/H_2_O: Hydro-methanolic extract of *Pinus sp.* leaves (30:70, v/v); Sodium diclofenac: Reference drug.

As shown in Figure 3, lower concentrations of plant extracts (less than 20 µg/mL) decreased the levels of nitric oxide, whereas cell treatment with 100 µg/mL of Ps.FMeOH/H₂O (6.55 µM), Ps.FMeOH (4.10 µM), and Ps.FEtOH/H₂O (3.98 µM) extracts and amphotericin B increased the levels of nitric oxide compared to the untreated control (3.41 µM). By contrast, the nitric oxide level was increased by cell treatment with Ps.FEtOH extract at 100 µg/mL.

#### 3-1-3. BSA anti-denaturation assay

The BSA anti-denaturation test was used to assay the anti-inflammatory potential of the plant extracts. Table 2 summarises the half-maximal inhibitory concentrations (IC_50_s) of extracts and diclofenac upon BSA anti-denaturation assay. Among the extracts tested, Ps.FMeOH/H_2_O extract showed the lowest IC_50_ value (4.11 µg/mL) followed by Ps.FMeOH (22.53 µg/mL), Ps.FEtOH/H_2_O (26.71 µg/mL), and Ps.FEtOH (52.38 µg/mL) extracts. Sodium diclofenac, which was used as the positive control, exhibited IC_50_ value of 2.91 µg/mL.

**Table 2:** IC_50_ values of *Pinus sp.* extracts and sodium diclofenac upon the BSA anti-denaturation assay. **P ≤ 0.01, ****P ≤ 0.0001 (Dunnett’s test).; Values are significantly different, compared to the positive control diclofenac sodium; Ps.FEtOH: Ethanol extract of *Pinus sp*. leaves; Ps.FMeOH: Methanol extract of *Pinus sp.* leaves; Ps.FEtOH/H_2_O: Hydro-ethanolic extract of *Pinus sp.* leaves (30:70, v/v); Ps.FMeOH/H_2_O: Hydro-methanolic extract of *Pinus sp.* leaves (30:70, v/v); IC_50_: Concentration that inhibits 50% of BSA denaturation; SD: Standard deviation; PC: Positive control.

| Serial number | Samples | IC<sub>50</sub>±SD (µg/ mL) |
| --- | --- | --- |
| 1 | Ps.FEtOH | 52.38±0.02**** |
| 2 | Ps.FMeOH | 22.53±0.03**** |
| 3 | Ps.FEtOH/H<sub>2</sub>O | 26.71±0.36**** |
| 4 | Ps.FMeOH/H<sub>2</sub>O | 4.11±0.04** |
| 5 | Sodium diclofenac | 2.91±0.01 |

As already discussed, the IC_50_ values of the extracts varied from 4.11±0.04 to 52.38±0.02 µg/mL, the most potent being the Ps.FMeOH/H_2_O extract (IC_50_: 4.11 µg/mL), compared with the value obtained for diclofenac sodium (IC_50_: 2.91 µg/mL).

#### 3-1-4. Total phenolic and flavonoid contents

Figure 5 illustrates the total phenolic and flavonoid contents of the four *Pinus sp.* extracts. Noteworthy, the highest phenolic content was observed in the ethanol extract (43.66 mg GAE/g), whereas the hydroethanolic extract (Ps.FEtOH/H₂O) revealed highest flavonoid content (43.89 mg QE/g). According to the statistical analysis, Ps.FEtOH and Ps.FMeOH extracts did not show any difference in their phenolic and flavonoid contents (p>0.05).

**Figure 5:**
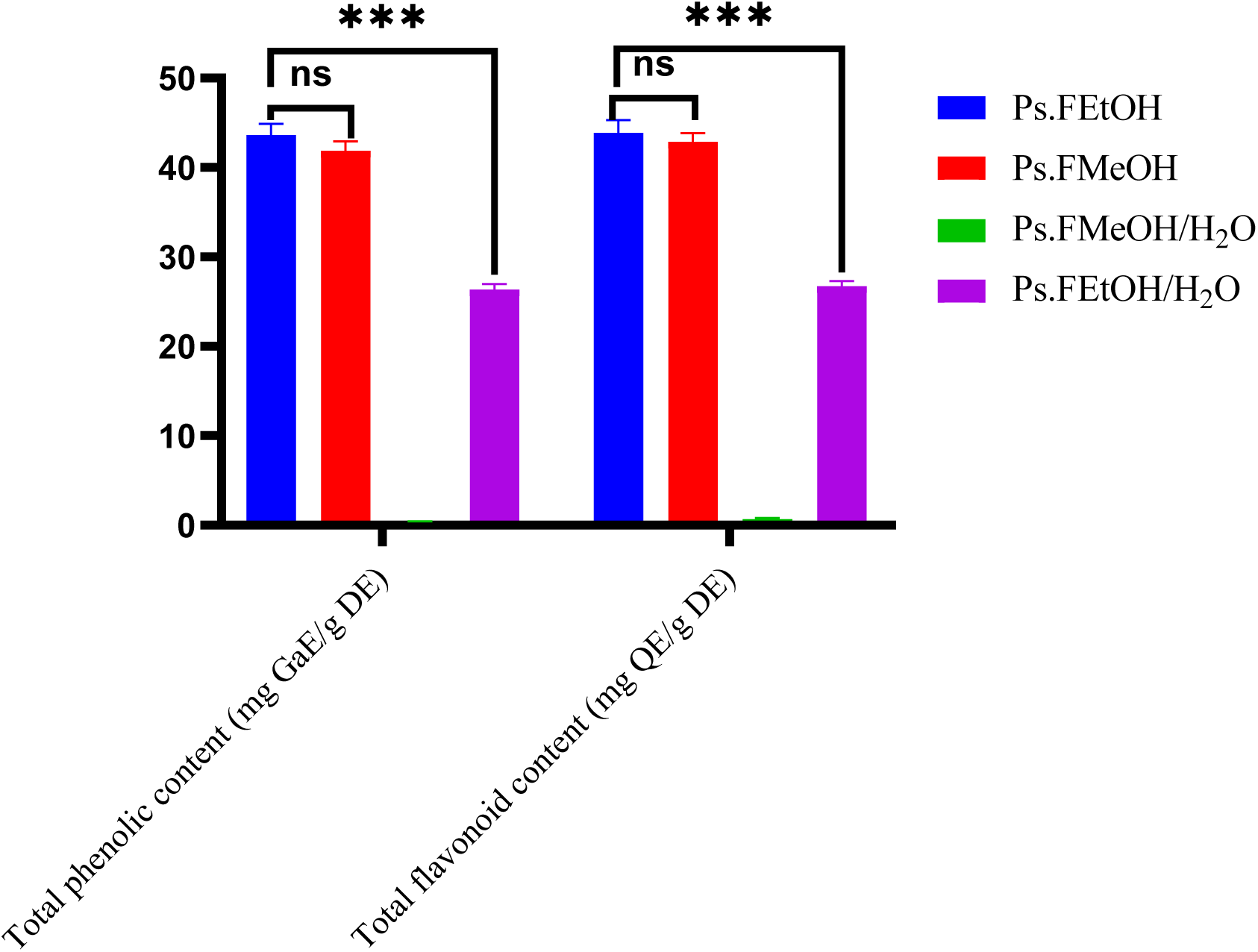
Total phenolic and flavonoid contents of extracts from *Pinus sp*. Leaves ***P ≤ 0.001, ns: Not significant (Dunnett’s test); Values are significantly different among the extract treatments; Ps.FEtOH: Ethanol extract of *Pinus sp*. leaves; Ps.FMeOH: Methanol extract of *Pinus sp*. leaves; Ps.FEtOH/H_2_O: Hydro-ethanolic extract of *Pinus sp*. leaves (30:70, v/v); Ps.FMeOH/H_2_O: Hydro-methanolic extract of *Pinus sp*. leaves (30:70, v/v); mg: mg: milligram; GaE/DE: gram equivalent of gallic acid per gram of dry extract; QE/DE: gram equivalent of quercetin per gram of dry extract.

#### 3-1-5. Antioxidant activity

The antioxidant effects of the *Pinus sp.* extracts were evaluated through DPPH, ABTS and FRAP assays. Table 3 summarizes median scavenging concentrations for DPPH and ABTS tests and 50% reduction of Fe^3+^ to Fe^2+^ for the FRAP assay.

**Table 3:** Median scavenging concentrations (SC_50_) and 50% reduction concentrations (RC_50_) of the *Pinus sp.* Extracts *P ≤ 0.05, **P ≤ 0.01, ***P˂0.001, ****(P˂0.0001), ns: Not significant (Dunnett’s test); Values are significantly different, compared to the value obtained for the positive control ascorbic acid; Ps.FEtOH: Ethanol extract of *Pinus sp.* leaves; Ps.FMeOH: Methanol extract of *Pinus sp.* leaves; Ps.FEtOH/H_2_O: Hydro-ethanolic extract of Pinus sp. leaves (30:70, v/v); Ps.FMeOH/H_2_O: Hydro-methanolic extract of Pinus sp. leaves (30:70, v/v); SC_50_: Concentration required to scavenge 50% of radicals; RC_50_: Concentration required to reduce 50% of Fe^3+^ to Fe^2+^ ; SD: Standard deviation; PC: Positive control; DPPH: 2,2-Diphenyl-1-picrylhydrazyl; ABTS: 2,2’-azinobis(3-ethylbenzothiazoline-6-sulfonic acid) diammonium salt; FRAP: Ferric reducing antioxidant power.

| Serial number | Samples | ABTS | DPPH | FRAP |
| --- | --- | --- | --- | --- |
| | | $SC_{50} \pm SD$ ($\mu\text{g/mL}$) | $SC_{50} \pm SD$ ($\mu\text{g/mL}$) | $RC_{50} \pm SD$ ($\mu\text{g/mL}$) |
| 1 | Ps.FEtOH | 205.1±2.31*** | 14.77±1.17* | 278.8±2.44*** |
| 2 | Ps.FMeOH | 144.4±2.16** | 8.98±0.96<sup>ns</sup> | 206.5±2.31*** |
| 3 | Ps.FEtOH/H<sub>2</sub>O | 88.16±1.95* | 22.8±1.36** | 111.2±2.04* |
| 4 | Ps.FMeOH/H<sub>2</sub>O | 81.14±1.90* | 25.12±1.40** | 359.7±2.55**** |
| 5 | Ascorbic acid | 0.61±0.21 | 6.85±0.84 | NT |
| 6 | Gallic acid | NT | NT | 1.60±0.20 |

The percentages of radicals scavenged following ABTS and DPPH tests, as well as the percentages of reduction of ferric (Fe^3+^) to ferrous (Fe^2+^) ions were also examined (Figure 6).

**Figure 6:**
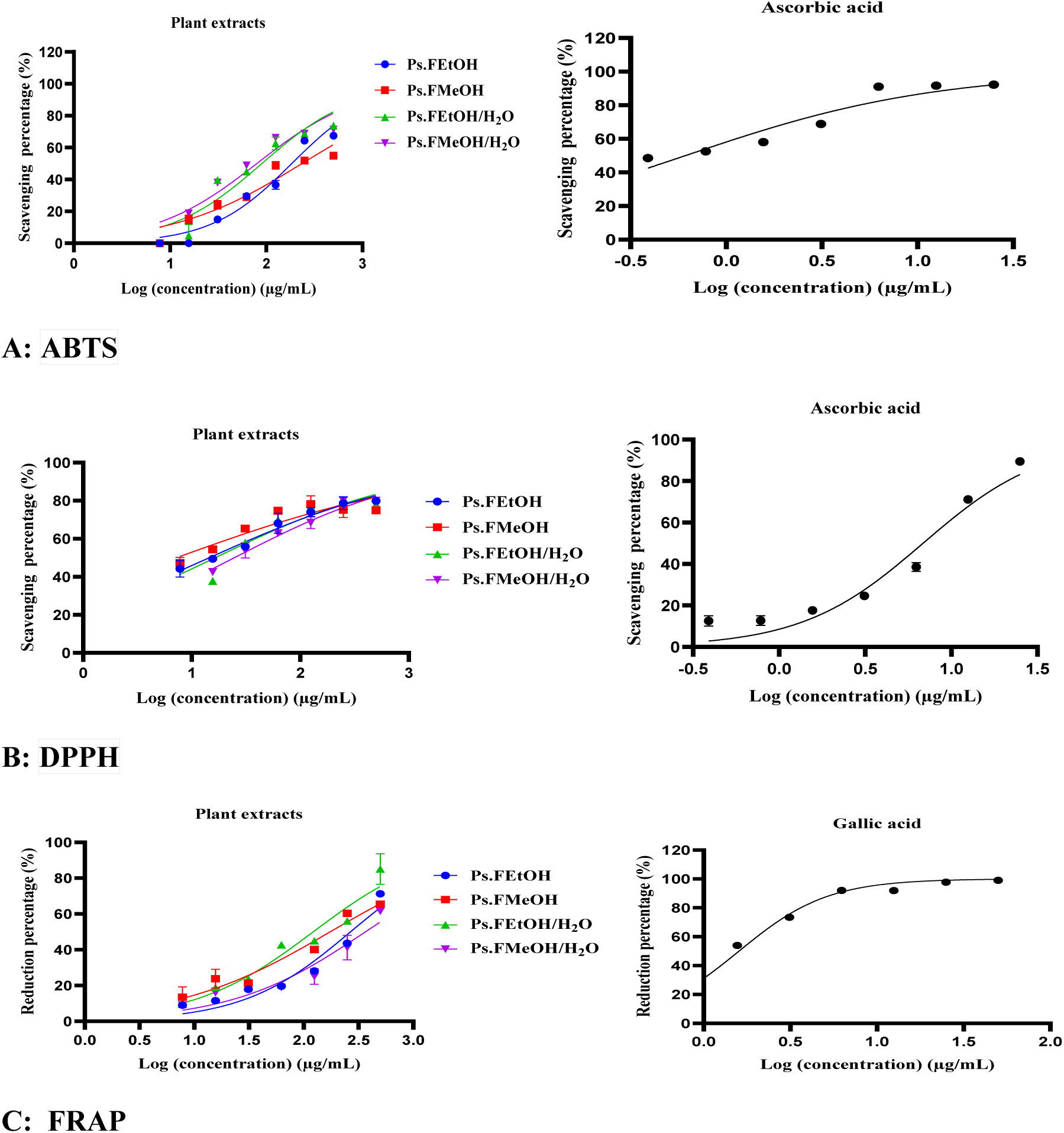
Percentages of radicals scavenged upon DPPH (A) and ABTS (B) tests and reduction percentages of ferric (Fe3+) to ferrous (Fe2+) ions (C) following treatment with Pinus sp. extracts and ascorbic acid.

The SC_50_ values ranged from 81.14 ± 1.90 to 205.1 ± 2.31 µg/mL, and 6.85 ± 0.84 to 25.12 ± 1.40 µg/mL, following ABTS and DPPH tests, respectively, whereas the RS_50_ values varied from 111.2 ± 2.0 to 359.7 ± 2.55 µg/mL upon the FRAP assay. In DPPH assay, the highest antiradical scavenging activity was observed with the Ps.MeOH extract, with a SC_50_ value close to that of ascorbic acid (SC_50_ : 6.85 ± 0.84 µg/mL), since there was no significant difference (P > 0.05) between these values.

#### 3-1-6. Acute toxicity

The most active antileishmanial was evaluated for oral toxicity in albino Wistar rats as per the OECD guidelines. As a result, there were no signs of toxicity up to 5000 mg/kg following the 14-days observation period, thus revealing an LD_50_ more than 5000 mg/kg.

##### 3-1-6-1. Effects of Ps.FEtOH extract on the morbidity of rats

The oral administration of Ps.FEtOH extract at 2000 and 5000 mg/kg did not induce any visible signs of toxicity (aggressiveness, mobility, piloerection, fecal aspect, etc.)in the experimental animals (Table 4). Moreover, there was no mortality recorded in treated rats at 2000 and 5000 mg/kg.

**Table 4:** Mortality and behavioural changes in rats following a single oral administration of the ethanol extract of *Pinus sp.* leaves after 14 days’ observation period. Ps.FEtOH: Ethanol extract of *Pinus sp*. leaves; N = Normal; N3 = Normal in all 3 animals; A = Absent; A3 = Absent in all 3 animals.

| Parameters | Control group<br>(10% Tween 80) | Ps.FEtOH@2000<br>mg/kg | Ps.FEtOH@5000<br>mg/kg |
| --- | --- | --- | --- |
| Number of deaths | 0/3 | 0/3 | 0/3 |
| Shivering / Chills | A3 | A3 | A3 |
| Aggressiveness | 0/3 | 0/3 | 0/3 |
| Mobility | 3/3 | 3/3 | 3/3 |
| Appearance of feces | N3 | N3 | N3 |
| Piloerection | 0/3 | 0/3 | 0/3 |
| Sensitivity to touch | 3/3 | 3/3 | 3/3 |
| Sensitivity to noise | 3/3 | 3/3 | 3/3 |

##### 3-1-6-2. Changes in rat body weights

During the experimental period (14 days), the animals were weighed every other day. Figure 7 illustrates the evolution of the rat body weights during the 14 days observation period. Treatment of rats with a single oral dose of the ethanol extract of *Pinus sp.* leaves induced a progressive increase in the rats’ body weights, when compared with the untreated animals.

**Figure 7:**
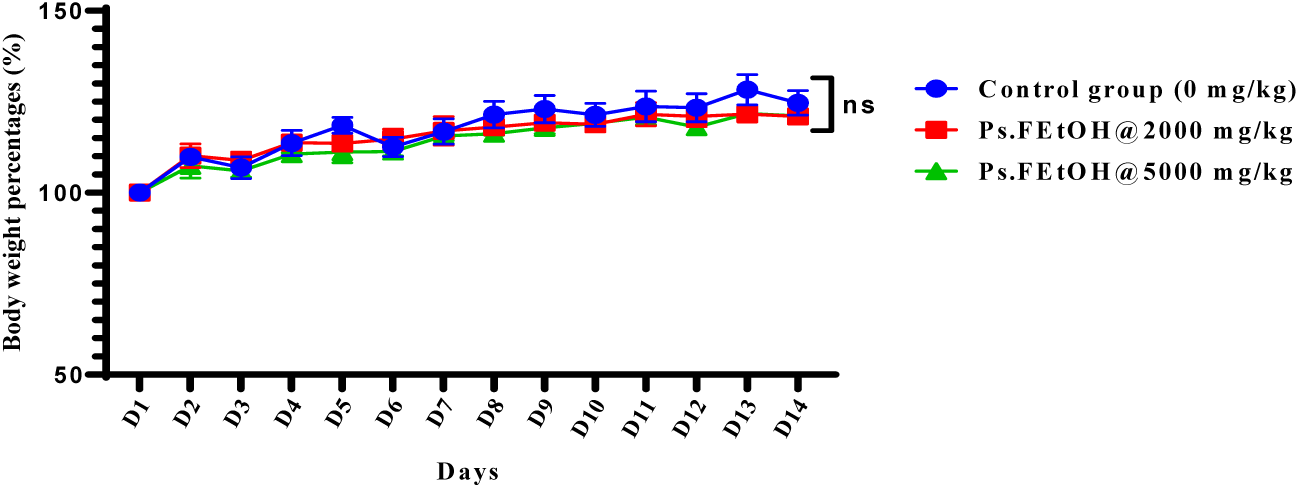
Effects of the ethanol extract of *Pinus sp.* leaves on the rat body weights ns: Not significant (Dunnett’s test); Values are significantly different, compared to the value obtained for the vehicle control group.

There was no significant difference (P>0.05) in the body weights of treated and untreated animals. As already discussed, the no observed adverse effect level (NOAEL) was observed at 5000 mg/kg, thus revealing a LD_50_ greater than 5000 mg/kg.

##### 3-1-6-3. Relative organ weights

After 14 days of observation, the single oral administration of the ethanol extract of *Pinus sp.* leaves at 2000 and 5000 mg/kg did not change the organ weights of the animals as compared with the untreated control rats (Figure 8).

**Figure 8:**
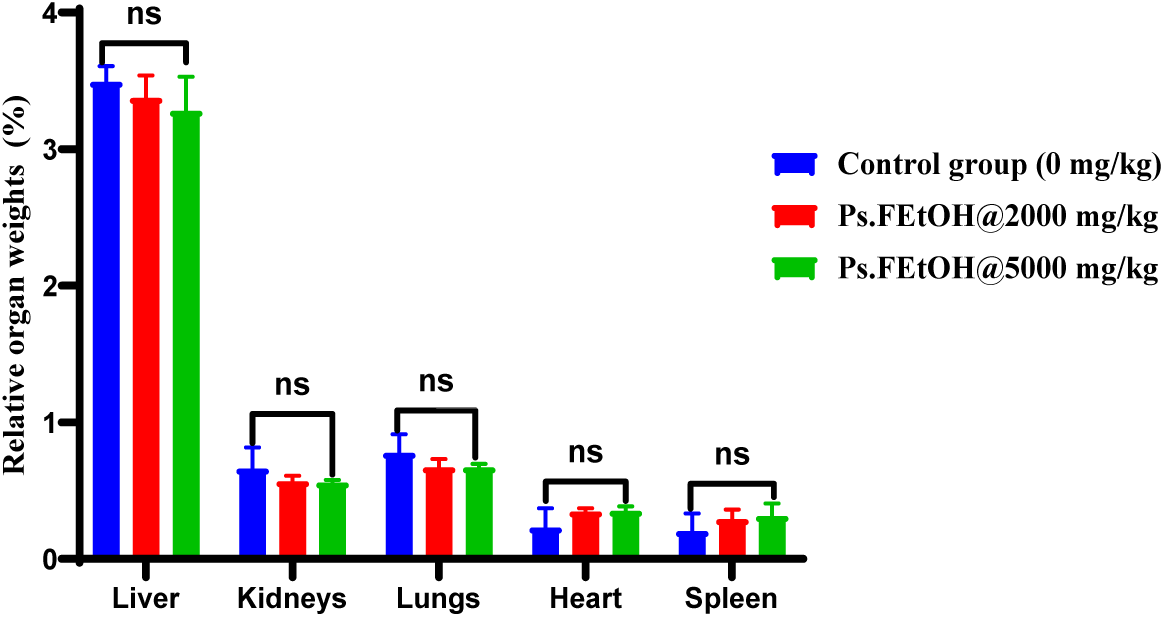
Relative organ weights treated with different extracts from *Pinus sp*. leaves, versus untreated animals. ns: Not significant (Dunnett’s test); Values are significantly different, compared to the value obtained for the vehicle control group.

Although there were no significant changes in the weights of organs, a notable increase in the weight of heart and spleen was observed, vs untreated group of animals.

##### 3-1-6-4. Histological sections of liver of rats

Figure 9 shows the effects of ethanol extract treatment on liver histology following the acute toxicity test. Microphotographs of liver sections from treated (Ps.FEtOH extract at 2000 and 5000 mg/kg) and untreated (10% Tween 80 in distilled water) animals display a well-structured portal vein (PV), normal hepatic parenchyma with hepatocytes (He) separated by sinusoidal capillaries (SC), and a visible hepatic artery (HA) and bile duct (BC). Notably, the hepatic parenchyma was well-organized on the liver histological sections of rats administered with the Ps.FEtOH extract as compared to the untreated control group (Figure 9).

**Figure 9:**
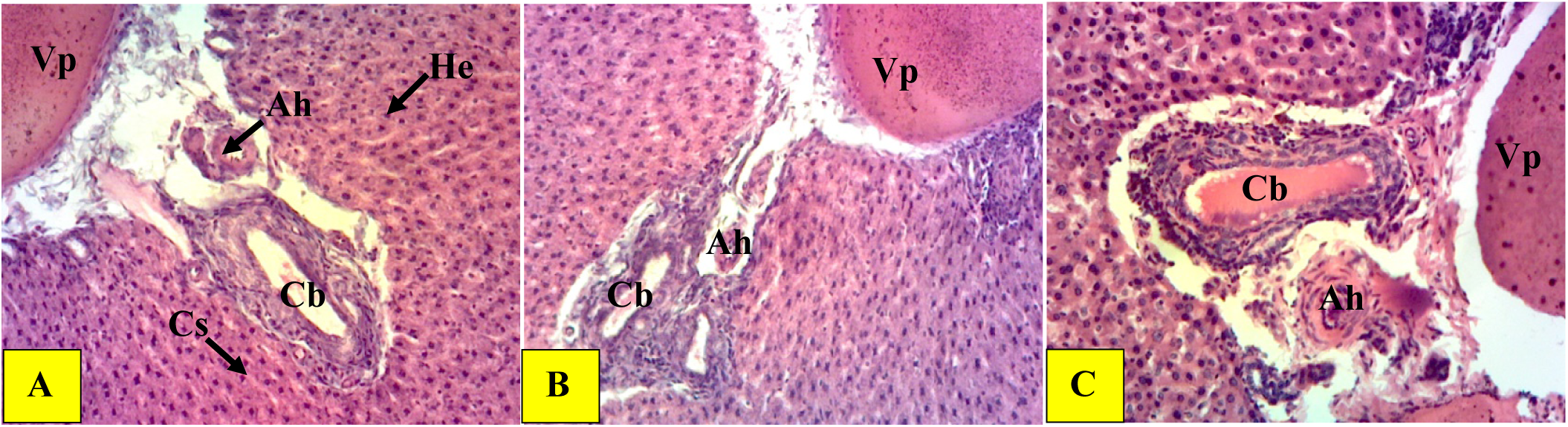
Histological section of the liver (200X, hematoxylin-eosin) of rats administered with single oral doses of Ps.FEtOH extract Vp = portal vein; Cb = bile duct; He = hepatocyte; Cs = sinusoidal capillary; Ah = hepatic artery; DCs = dilation of sinusoidal capillaries; Fi = inflammatory focus; A = Normal control, B = Ps.FEtOH2000, C = Ps.FEtOH5000.

##### 3-1-6-5. Effects of ethanol extract on rat kidney histology

The microarchitecture of the kidneys of treated (@2000 and 5000 mg/kg of Ps.FEtOH extract) and untreated (10% Tween 80 in distilled water) animals shows a typical renal parenchyma, in which the glomerulus (Gl), Bowman’s space or urinary space (BS), and the proximal (PT) and distal convoluted tubules (DT) are normal and well-differentiated. The parenchymal architecture of the kidneys of treated and untreated rats appeared to be normal (Figure 10).

**Figure 10:**
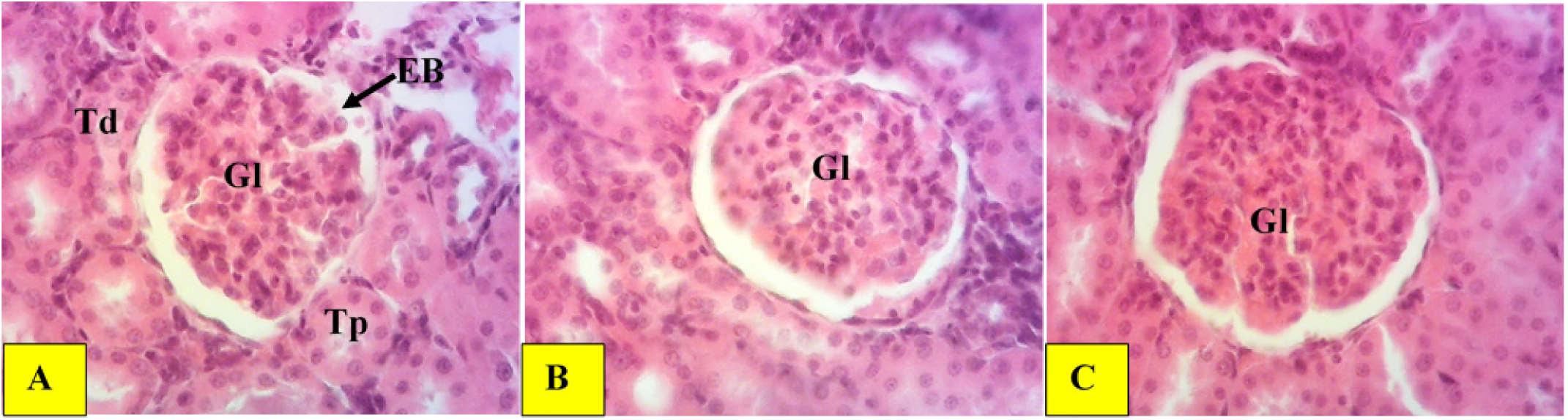
Histological section of the kidney (200X, hematoxylin-eosin) of rats administered with single oral doses of Ps.FEtOH extract Gl = Glomerulus; Td = Distal convoluted tubule; Tp = Proximal convoluted tubule; BS = Bowman’s space; A = Normal control; B = Group of rats administered with the ethanol extract at 2000 mg/kg ; C = Group of rats treated with the ethanol extract at 5000 mg/kg.

##### 3-1-6-6. Hematological parameters

The single oral administration of the ethanol extract of *Pinus sp.* leaves at 2000 and 5000 mg/kg did not induce any significant changes (P>0.05) in the haematological parameters of treated rats (Table 5).

**Table 5:** Effects of the ethanol extract on the hematological parameters in rats. a: Values are means±SD of three animals in each group (n = 3); ns: Not significant (Dunnett’s test); Values are significantly different, compared to that obtained for the vehicle control; RBC: Red blood cells; HCT: Hematocrit; HGB: Hemoglobin concentration; WBC: White Blood Cells; PLT: Platelets; MCV: Mean Corpuscular Volume; MCH: Mean corpuscular haemoglobin; MCHC: Mean corpuscular haemoglobin concentration; LYM: Lymphocytes; MON: Monocytes; GRA: Granulocytes; Ps.FEtOH@2000: 2000 mg/kg of ethanol extract from *Pinus sp.* leaves; Ps.FEtOH@5000: 5000 mg/kg of ethanol extract from *Pinus sp.* leaves; VC: Vehicle control (group of animals that received 10% Tween 80 diluted in distilled water).

| Parameters<sup>a</sup> | Control group<br>(VC) | PsFETOH@2000 | PsFETOH@5000 |
| --- | --- | --- | --- |
| WBC ($10^3/\text{mm}^3$) | 8.3±0.45 | 9.1±1.6<sup>ns</sup> | 9.6±1.17<sup>ns</sup> |
| RBC ($10^6/\text{mm}^3$) | 9.27±0.53 | 8.67±0.51<sup>ns</sup> | 9.4±0.53<sup>ns</sup> |
| HGB (g/dL) | 16.36±1.12 | 14.8±1.5<sup>ns</sup> | 17.3±1.66<sup>ns</sup> |
| HCT (%) | 45.06±5.11 | 40.6±1.7<sup>ns</sup> | 45.1±5.1<sup>ns</sup> |
| PLT ($10^3/\text{mm}^3$) | 359.66±14.57 | 337±19.69<sup>ns</sup> | 351.33±20.3<sup>ns</sup> |
| MCV (fl) | 90.33±1.52 | 90.66±4.72<sup>ns</sup> | 98.66±8.5<sup>ns</sup> |
| MCH (pg) | 39.1±1.11 | 34.06±3.88<sup>ns</sup> | 37.6±1.8<sup>ns</sup> |
| MCHC(g/dL) | 41.4±1.82 | 38.03±6<sup>ns</sup> | 40.83±4.34<sup>ns</sup> |
| LYM (%) | 32.3±2.57 | 36.46±8.71<sup>ns</sup> | 39.5±9.02<sup>ns</sup> |
| MON (%) | 4.36±2.76 | 6.86±1.3<sup>ns</sup> | 5.76±1.05<sup>ns</sup> |
| GRA (%) | 48.03±5.85 | 54.16±3.66<sup>ns</sup> | 66.06±13.82<sup>ns</sup> |
| LYM ($10^3/\text{mm}^3$) | 2.4±1.51 | 3.43±1.41<sup>ns</sup> | 3.73±0.06<sup>ns</sup> |
| MON ($10^3/\text{mm}^3$) | 0.56±0.15 | 1.3±1.05<sup>ns</sup> | 1.13±0.15<sup>ns</sup> |
| GRA ($10^3/\text{mm}^3$) | 3.56±1.6 | 3.26±0.32<sup>ns</sup> | 3.93±0.56<sup>ns</sup> |

##### 3-1-6-7. Biochemical parameters of toxicity

The single oral treatment of rats with the ethanol extract at 2000 and 5000 mg/kg did not induce any significant changes on liver biochemical parameters of toxicity, including ALT, AST and ALP (Table 6). There were no significant changes in the toxicity parameters of kidneys, such as creatinine and uric acid, even though the treatment of rats with 5000 mg/kg ethanol extract showed elevated values (1.83 and 5.06 mg/dL) compared with the vehicle control (1.79 and 3.91 mg/dL).

**Table 6:** Biochemical markers of toxicity in liver and kidneys of rats administered with the plant extracts a: Values are means±SD of three animals in each group (n = 3); ns: Not significant (Dunnett’s test); Values are significantly different, compared to that obtained for the vehicle control; IU: International unit; ALT: Alanine aminotransferase; AST: Aspartate aminotransferase; ALP: Alkaline phosphatase; Ps.FEtOH@2000: 2000 mg/kg ethanol extract of *Pinus sp.* leaves; Ps.FEtOH@5000: 5000 mg/kg ethanol extract of *Pinus sp.* leaves; Control group (vehicle control): Group of animals that received only 10% Tween 80 diluted in distilled water.

| Biochemical parameters | Control group<br>(vehicle control) | Ps.FEtOH@2000 | Ps.FEtOH@5000 |
| --- | --- | --- | --- |
| Kidneys |  |  |  |
| **Creatinine (mg/dL)** | 1.79±0.15 | 1.75±0.04<sup>ns</sup> | 1.83±0.1<sup>ns</sup> |
| **Uric acid (mg/dL)** | 3.91±0.58 | 4.63±1.12<sup>ns</sup> | 5.06±0.55<sup>ns</sup> |
| **Protein (mg/mL)** | 0.25±0.03 | 0.25±0.04<sup>ns</sup> | 0.26±0.03<sup>ns</sup> |
| Liver |  |  |  |
| **AST (UI)** | 114.59±15.98 | 116.3±10.51<sup>ns</sup> | 112.02±6.53<sup>ns</sup> |
| **ALT (UI)** | 44.31±6.23 | 51.31±11.35<sup>ns</sup> | 47.8±9.88<sup>ns</sup> |
| **ALP (UI/L)** | 103.38±13.58 | 110.91±12.25<sup>ns</sup> | 116.52±22.36<sup>ns</sup> |

#### 3-1-7. Phytochemical analysis

The chemical profile of the most active extract, Ps.FEtOH, was analyzed by UHPLC-LC-MS/MS to identify the compounds likely responsible for its observed biological activities (antileishmanial, antioxidant and anti-inflammatory activities). From the raw data, a total of 38 mass features were detected. Preliminary identification using various databases yielded 20 potential compounds. Among these, 9 compounds were selected based on a high matching score (>0.90) from the mass spectrometry library, thus ensuring a high degree of confidence in the structural identification (Table 7). Several families of compounds were identified in Ps.FEtOH extract, including dicarboxylic acids and derivatives; p-methoxybenzoic acids and derivatives; alkyl-phenylketones; alkaloids; 7-O-methylated flavonoids; colensane and clerodane diterpenoids; long-chain fatty acids; and diterpenoids (Table 7). Figure 11 illustrates the chemical structures of annotated compounds of the Ps.FEtOH extract.

**Figure 11:**
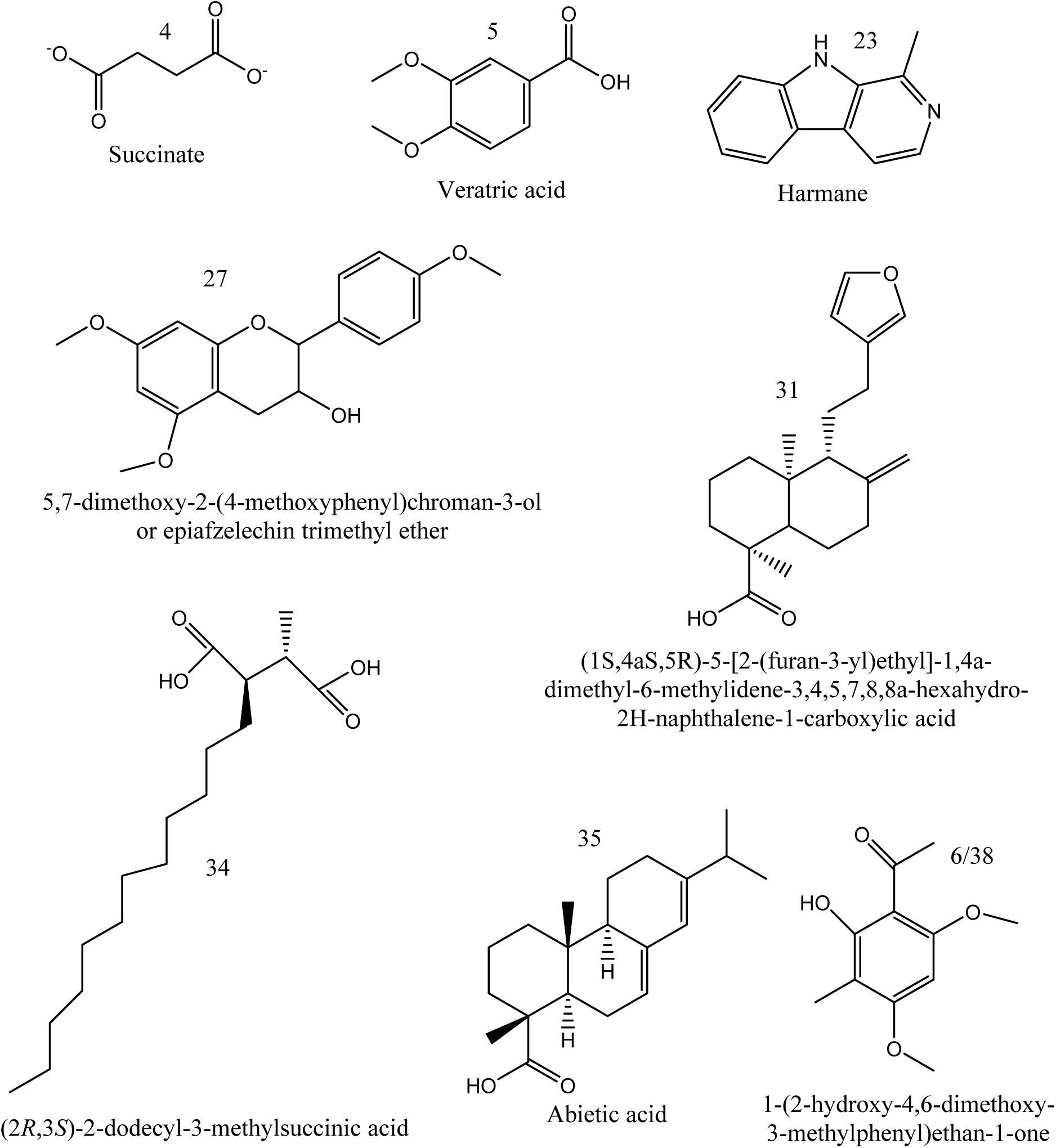
Chemical structures of compounds annotated in the Ps.FEtOH extract.

**Table 7:** Secondary metabolites annotated in the PsEtOH extract using UHPLC-LCMS/MS in negative ionization mode Mass spectra were acquired in a QTOF mass spectrometer equipped with a HESI source operating in negative ion mode; RT: Retention time; m/z: mass on charge; [M-H]: compound minus 1 M mass; Adduct: ionization mode (negative).

| ID | Name of the metabolite | Family of compounds | Molecular formula | m/z | RT (min) | Adduct | Matching score |
| --- | --- | --- | --- | --- | --- | --- | --- |
| 4 | Succinate | Dicarboxylic acids and derivatives | C<sub>4</sub>H<sub>6</sub>O<sub>4</sub> | 112.98924 | 0.0813167 | [M+HCOO]<sup>-</sup> | 0.9971516 |
| 5 | Veratric acid | P-methoxybenzoic acids and derivatives | C<sub>9</sub>H<sub>10</sub>O<sub>4</sub> | 180.981125 | 0.08131667 | [M+TFA-H]<sup>-</sup> | 0.9017829 |
| 6 | 1-(2-hydroxy-4,6-dimethoxy-3-methylphenyl)ethan-1-one/Methylxanthoxylin | Alkyl-phenylketones | C<sub>11</sub>H<sub>14</sub>O<sub>4</sub> | 248.969635 | 0.08131667 | [M+HCOO]<sup>-</sup> | 0.9916243 |
| 23 | Harmane | Alkaloids | C<sub>12</sub>H<sub>10</sub>N<sub>2</sub> | 181.077484 | 0.36803333 | [M-H]<sup>-</sup> | 0.9993297 |
| 27 | 5,7-dimethoxy-2-(4-methoxyphenyl)chroman-3-ol/Epiafzelechin Trimethyl Ether | 7-O-methylated flavonoids | C<sub>18</sub>H<sub>20</sub>O<sub>5</sub> | 315.201447 | 4.18715 | [M-H]<sup>-</sup> | 0.9706374 |
| 31 | (1S,4aS,5R)-5-[2-(furan-3-yl)ethyl]-1,4a-dimethyl-6-methylidene-3,4,5,7,8,8a-hexahydro-2H-naphthalene-1-carboxylic acid | Colensane and clerodane diterpenoids | C<sub>20</sub>H<sub>28</sub>O<sub>3</sub> | 315.202576 | 4.69301667 | [M-H]<sup>-</sup> | 1.558251 |
| 34 | (2R,3S)-2-dodecyl-3-methylsuccinic acid/Roccellic acid | Long-chain fatty acids | C<sub>17</sub>H<sub>32</sub>O<sub>4</sub> | 299.210724 | 5.8859 | [M-H]<sup>-</sup> | 1.486351 |
| 35 | Abietic acid | Diterpenoids | C<sub>20</sub>H<sub>30</sub>O<sub>2</sub> | 301.227203 | 6.5897 | [M-H]<sup>-</sup> | 1.613123 |
| 38 | 1-(2-hydroxy-4,6-dimethoxy-3-methylphenyl)ethan-1-one/Methylxanthoxylin | Alkyl-phenylketones | C<sub>11</sub>H<sub>14</sub>O<sub>4</sub> | 248.971176 | 9.01768333 | [M-H]<sup>-</sup> | 0.9605252 |

#### 3-1-8. Molecular docking and prediction of ADMET properties

The molecular interactions between selected annotated compounds (harmane, abietic acid and epiafzelechin trimethyl ether) of the Ps.FEtOH extract were studied with the target enzyme trypanothione reductase using Autodock Vina. The pharmacokinetic properties of these compounds were also predicted.

##### 3-1-8-1. Prediction of ADMET properties

From the 9 compounds identified by LC-MS/MS, harmane, abietic acid, and epiafzelechin trimethyl ether were selected for the prediction of the ADMET properties and the results are summarized in Table 8. These compounds were chosen due to their structural novelty and/or their previously reported antileishmanial activity (Di Giorgio. C, 2004; Torres, 2020), allowing for a direct comparison with the existing antileishmanial drugs amphotericin B and miltefosine.

**Table 8:** Predicted physicochemical, pharmacokinetic, and toxicity parameters HBA: Hydrogen bond acceptors; HBD: Hydrogen bond donors; QED: Quantitative estimate of druglikeness; PTSA: Topological polar surface area; DILI: Drug-induced liver injury; BBB: Blood-brain barrier.

| Parameters | Harmane | Epiefzelechin trimethyl ether | Abietic acid | Amphotericin B | Miltefosine |
| --- | --- | --- | --- | --- | --- |
| **Physicochemical properties** |  |  |  |  |  |
| Molecular weight (Dalton) | 182.23 | 316.35 | 302.46 | 924.09 | 407.58 |
| LogP (log-ratio) | 3.02 | 2.75 | 5.21 | 0.71 | 5.68 |
| HBA | 1 | 5 | 1 | 17 | 4 |
| HBD | 1 | 1 | 1 | 12 | 0 |
| Lipinski rule of 5 | 4 | 4 | 3 | 1 | 3 |
| QED | 0.57 | 0.94 | 0.76 | 0.17 | 0.15 |
| TPSA ($\text{\AA}^2$) | 28.68 | 57.15 | 37.3 | 319.61 | 58.59 |
| **Pharmacokinetics** |  |  |  |  |  |
| Human intestinal absorption | 1 | 1 | 1 | 0.03 | 0.2 |
| Oral bioavailability | 0.91 | 0.94 | 0.92 | 0.27 | 0.35 |
| Aqueous solubility (log(mol/L)) | -2.75 | -3.09 | -4.79 | -3.14 | -3.86 |
| BBB penetration | 0.97 | 0.47 | 0.68 | 0.06 | 0.38 |
| Half-life (hour) | 13.3 | 0 | 0 | 12.2 | 29.13 |
| Volume of distribution (L/kg) | 8.2 | 0.38 | 0.11 | 0 | 0.97 |
| **Toxicity profile** |  |  |  |  |  |
| hERG Blocking | 0.27 | 0.7 | 0.11 | 0.13 | 0.95 |
| Clinical toxicity | 0.04 | 0.02 | 0.07 | 0.05 | 0.0002 |
| Mutagenicity | 0.81 | 0.09 | 0.04 | 0.15 | 0.01 |
| DILI | 0.89 | 0.54 | 0.38 | 0.35 | 0.02 |
| Carcinogenicity | 0.04 | 0.04 | 0.21 | 0.02 | 0.19 |

HBA: Hydrogen bond acceptors; HBD: Hydrogen bond donors; QED: Quantitative estimate of druglikeness; TPSA: Topological polar surface area; DILI: Drug-induced liver injury; BBB: Blood-brain barrier.

Study of the ADMET properties of harmane, epiafzelechin trimethyl ether, and abietic acid revealed that these compounds comply with Lipinski’s Rule of 5. Harmane and epiafzelechin trimethyl ether did not violate any of the Lipinski rules, whereas abietic acid violates only 1 rule. By contrast, amphotericin B violated 3 of the 5 rules, whereas miltefosine violated 2 rules. These results demonstrate that the test compounds have more favorable physicochemical properties (low molecular weight, low TPSA and high QED) than the reference drugs miltefosine and amphotericin B (low solubility, high TPSA, and low QED). The prediction of the pharmacokinetic properties revealed a high intestinal absorption (absorption index: 1) and an elevated oral bioavailability (>0.9) of the test compounds harmane, epiafzelechin trimethyl ether, and abietic acid when compared with the reference drugs miltefosine (absorption and bioavailability indices: 0.2 and 0.27) and amphotericin B (absorption and bioavailability indices: 0.03 and 0.35).

Harmane is more likely to cross the blood-brain barrier (0.97) than the other test compounds and the reference antileishmanial drugs miltefosine (0.38) and amphotericin B (0.06). This compound also showed a half-life (13.3 h) comparable to that of amphotericin B (12.2); however, the prediction of its mutagenic (high mutagenic index: 0.81) and hepatotoxic (DILI: 0.89) riks did not show encouraging results. The high indices of hERG blocking (0.95) and skin reactions (0.95) predicted for miltefosine suggests potential heart and skin toxicity for this drug. Interestingly, epiafzelechin trimethyl ether revealed the lowest score for clinical toxicity 0.02) than the other test compounds.

Figure 12 is a radar chart that illustrates in percentage, the solubility, bioavailability, BBB safety, non-toxicity, and hERG safety. According to Figure 12, epiafzelechin trimethyl ether and abietic acid revealed high score (75-100%) for solubility and bioavailability; amphotericin B showed poor BBB properties, whereas harmane revealed favorable ADME properties but elevated toxicity.

**Figure 12:**
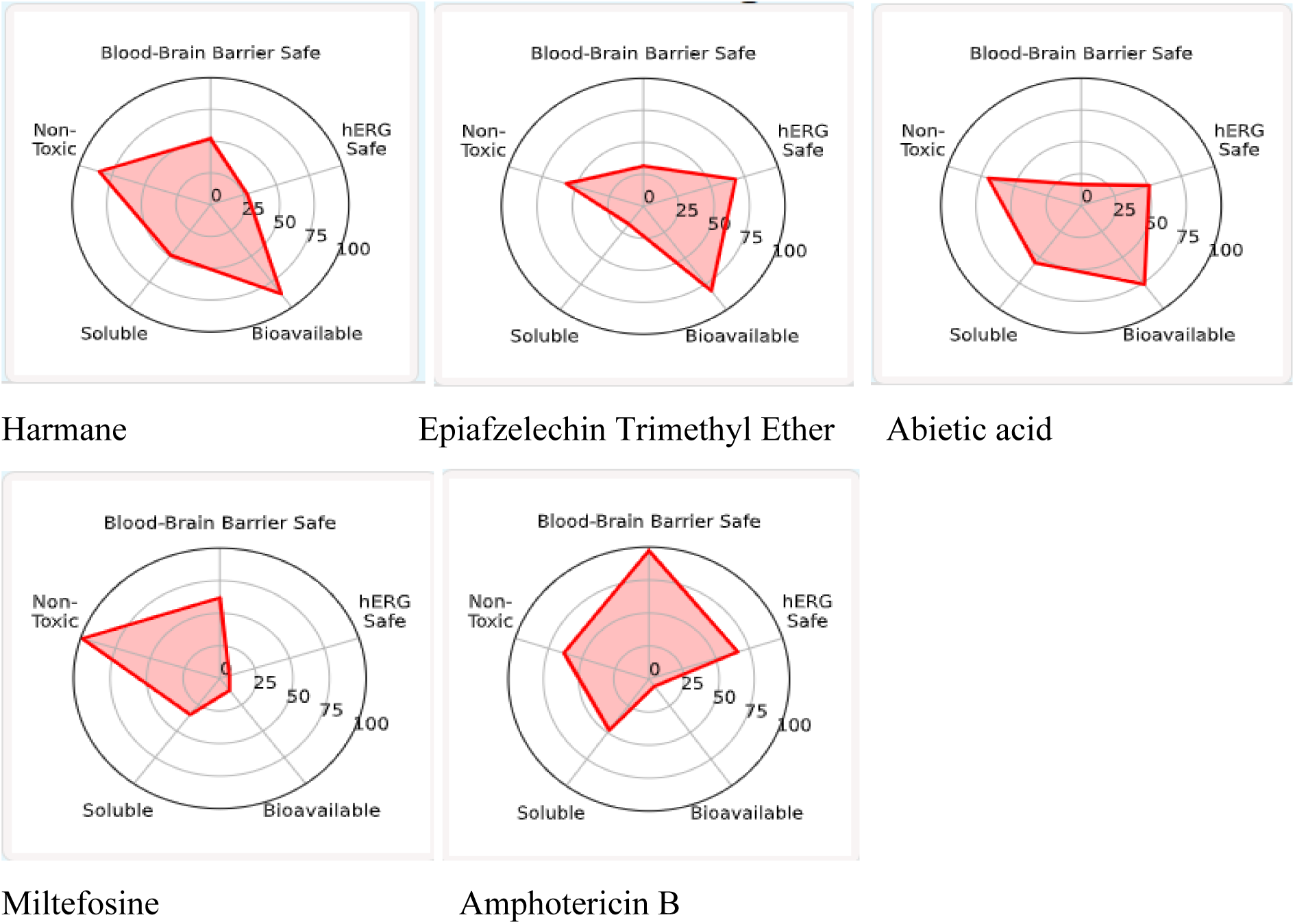
Radar chart showing the ADMET properties of the test compounds and the reference drugs miltefosine and amphotericin B.

The toxicity profile of the test compounds was also predicted using the DrugBank database. Figure 13 is a bar graph showing the predicted toxicity profiling of the test compounds, versu miltefosine and amphotericin. According to Figure 13, the test compounds revealed mild clinical toxicity and high intestinal absorption profile (score_∼_1).

**Figure 13:**
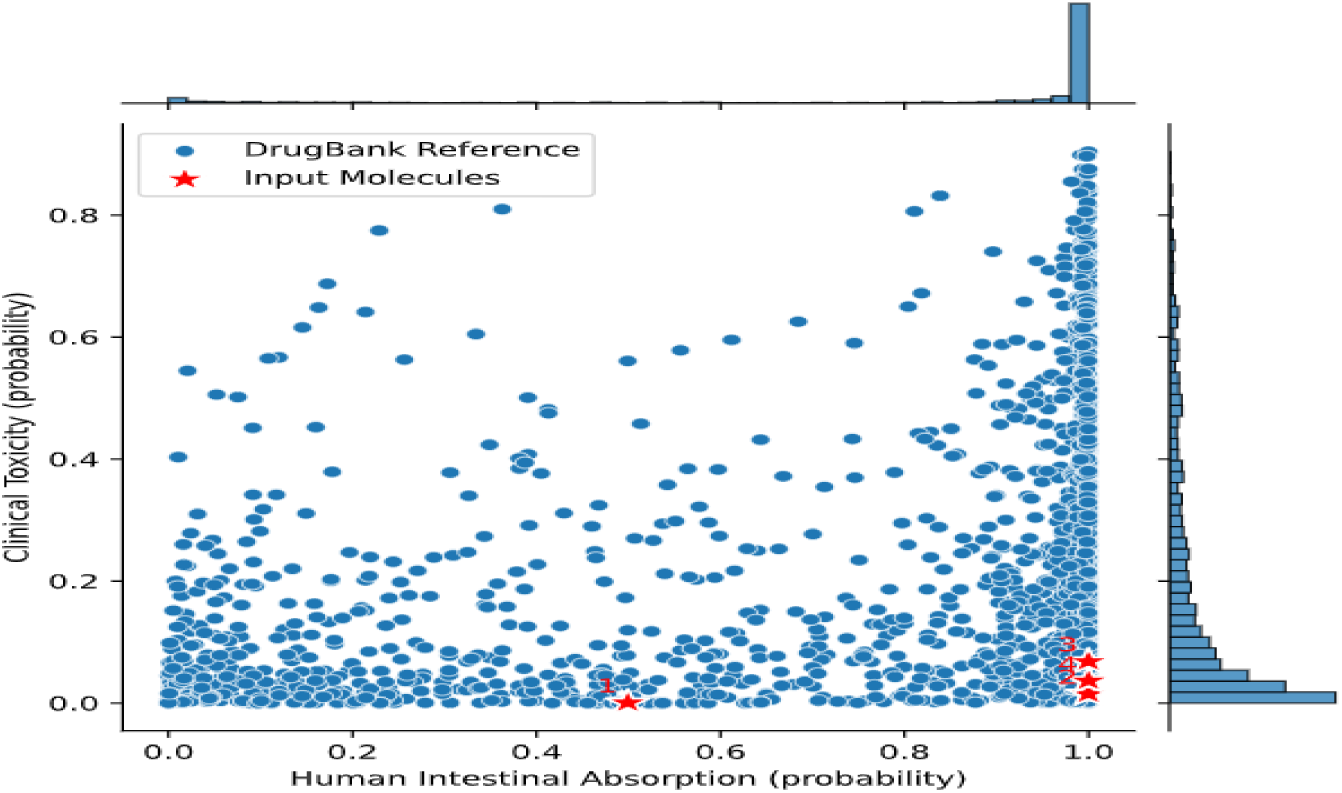
Bar graph showing the toxicity profile of the test compounds and reference drugs upon prediction using the DrugBank database.

##### 3-1-8-2. Molecular docking

The ligand-receptor interactions were predicted using Autodock Vina. Table 9 summarises the binding energies obtained with different complexes formed individually between the test compounds (ligands) and the target enzyme trypanothione reductase. Amphotericin B and miltefosine were included as the reference antileishmanial drugs. The binding energies of the test compounds ranged from −6.9 to −8.2 Kcal/moL, versus amphotericin B (−6.1 Kcal/moL) and miltefosine (−8.1 Kcal/moL). The lowest binding energy was recorded with harmane (−8.2 Kcal/moL), whereas abietic acid revealed the highest binding energy (−6.9 Kcal/moL). These results suggest that harmane displays highest binding affinity, whereas abietic acid exhibits the lowest binding affinity vis-à-vis trypanothione reductase, the target enzyme.

**Table 9:**
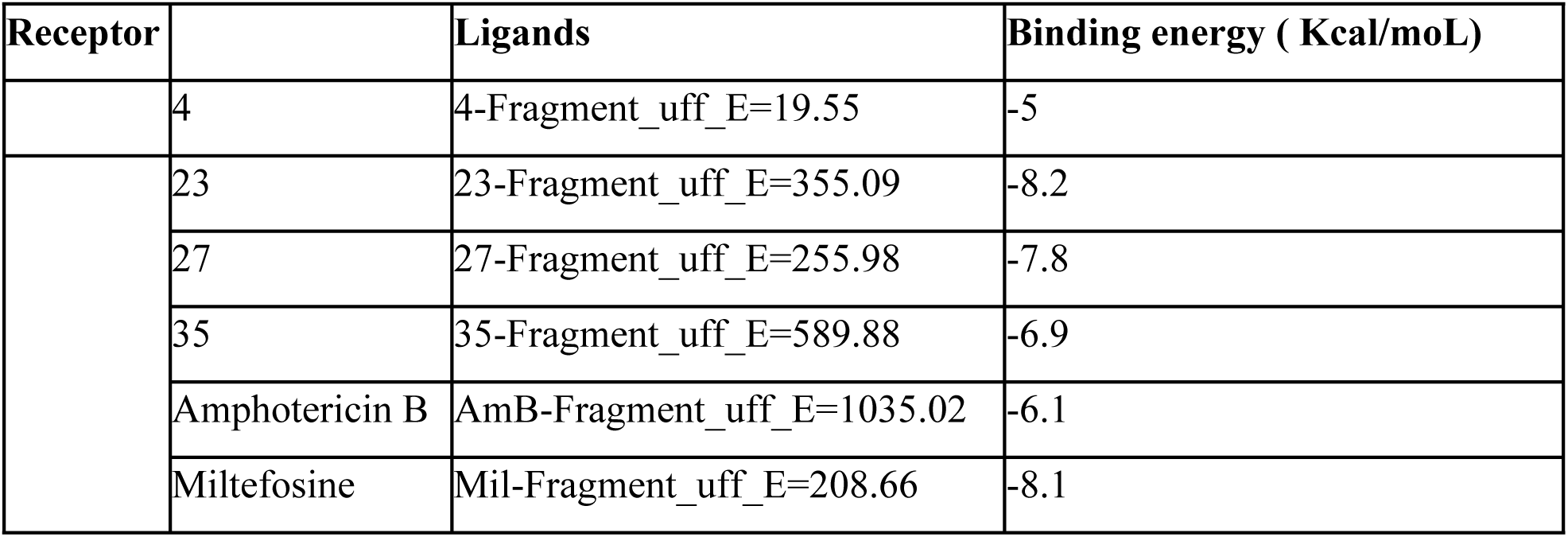
Binding energy (Kcal/moL) of the ligand-receptor complexes.

| Receptor |  | Ligands | Binding energy ( Kcal/mol) |
| --- | --- | --- | --- |
|  | 4 | 4-Fragment_uff_E=19.55 | -5 |
|  | 23 | 23-Fragment_uff_E=355.09 | -8.2 |
|  | 27 | 27-Fragment_uff_E=255.98 | -7.8 |
|  | 35 | 35-Fragment_uff_E=589.88 | -6.9 |
|  | Amphotericin B | AmB-Fragment_uff_E=1035.02 | -6.1 |
|  | Miltefosine | Mil-Fragment_uff_E=208.66 | -8.1 |

The molecular interactions between the test compounds and trypanothione reductase indicated the presence of a number of amino acid residues in the binding pocket. Table 10 summarizes the amino acid residues and the type of interactions at the binding site.

**Table 10:**
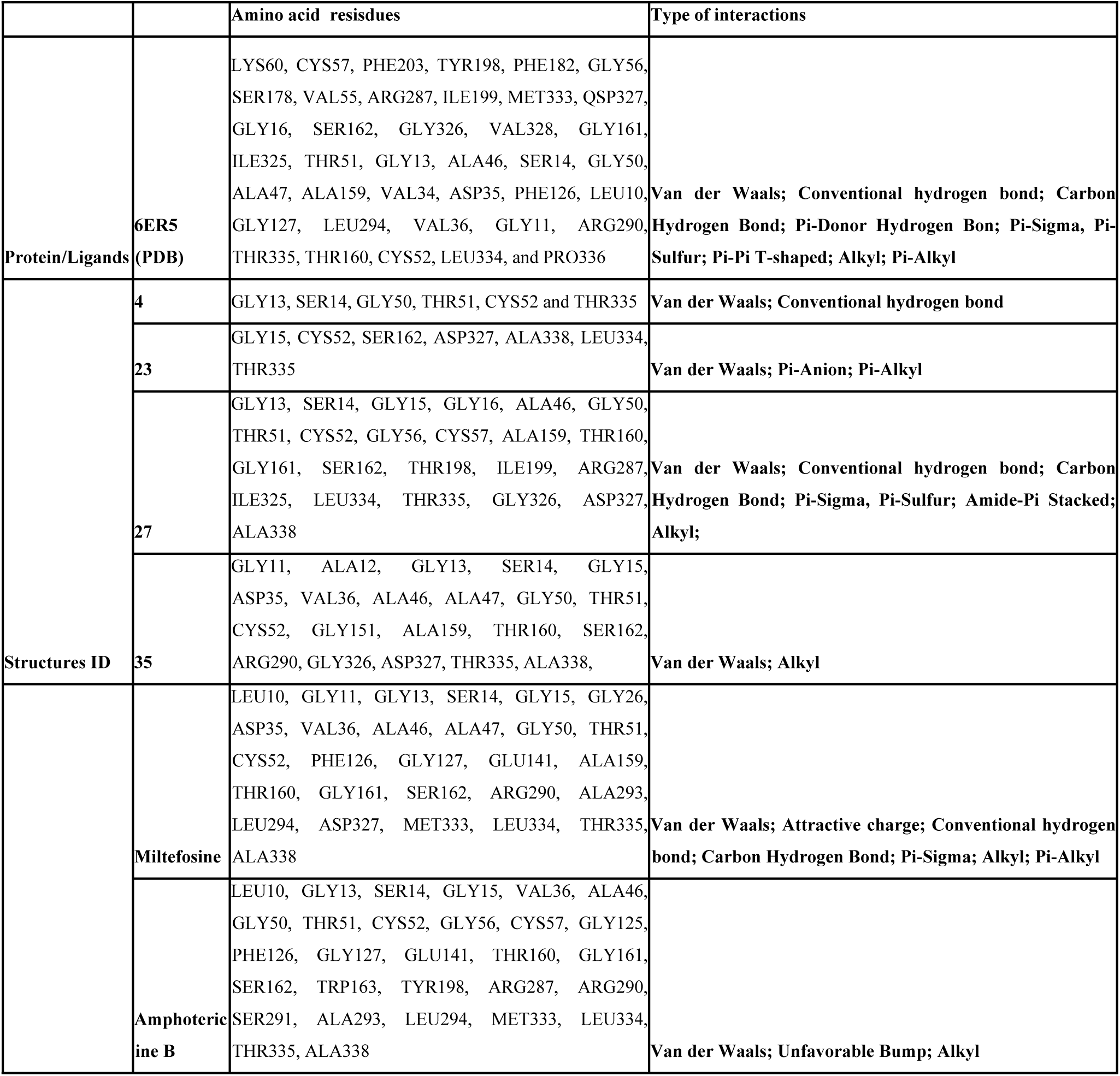
Amino acid residues and type of interactions recorded at the binding site of the ligand-receptor complexes.

Noteworthy, amino acids, such as GLY15, CYS52, SER162, ASP327, ALA338, LEU334 and THR335 were found in the binding pocket of the harmane-trypanothione reductase complex. These amino-acids were also found in the binding pockets of abietic acid and epiafzelechin trimethyl ether with trypanothione reductase (Table 10, Figure 14). Common molecular interations included Van der Waals, Pi-Anion and Pi-Alkyl interactions and hydrogen bonds (Figures 14 & 15).

**Figure 14.**
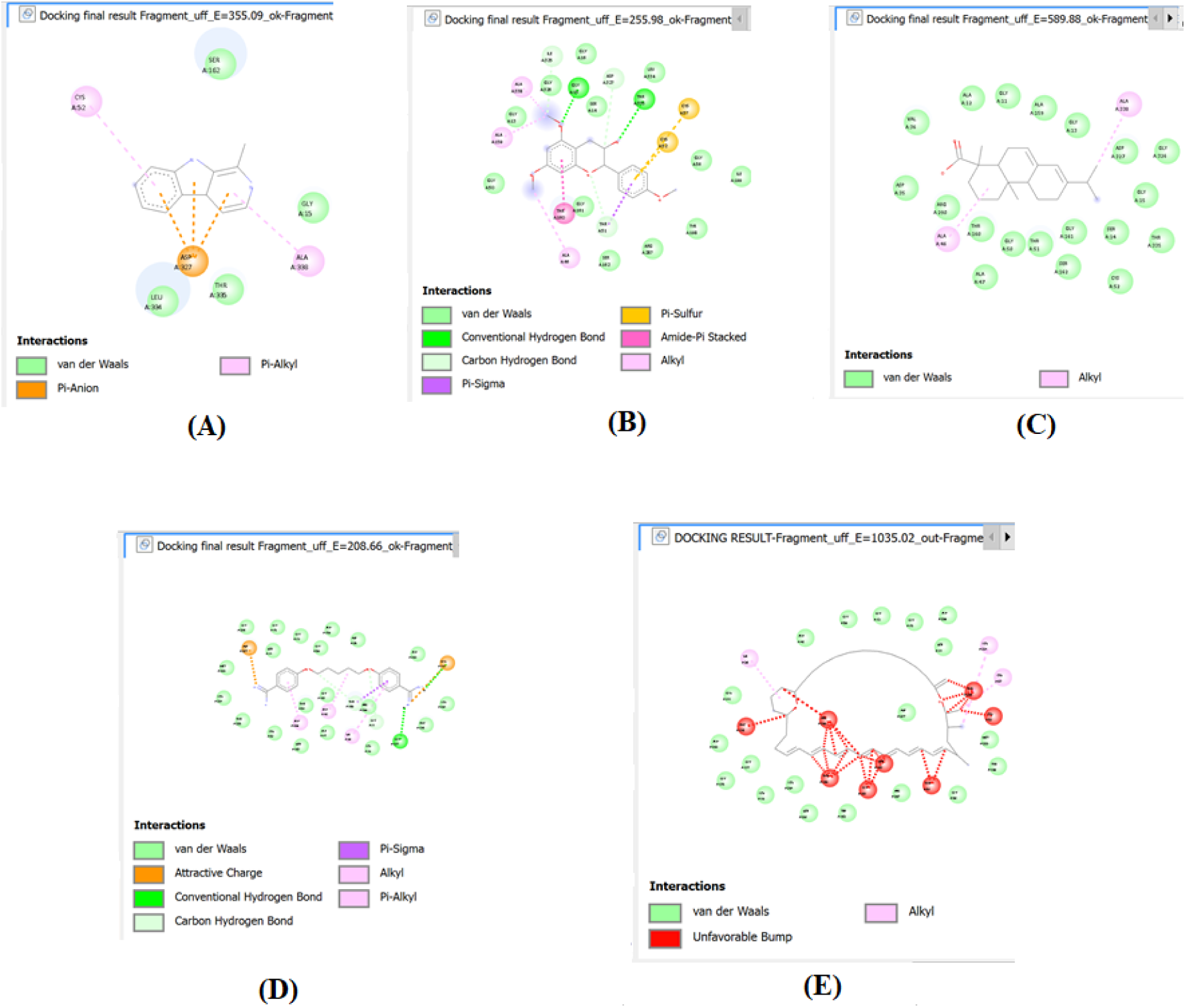
Protein-ligand interactions profiles. A, B, C, D and E corresponds respectively to the following complexes : harmane-6ER5, abietic acid-6ER5, epiafzelechin trimethyl ether-6ER5, miltefosine-6ER5 and amphotericin-6ER5.

**Figure 15:**
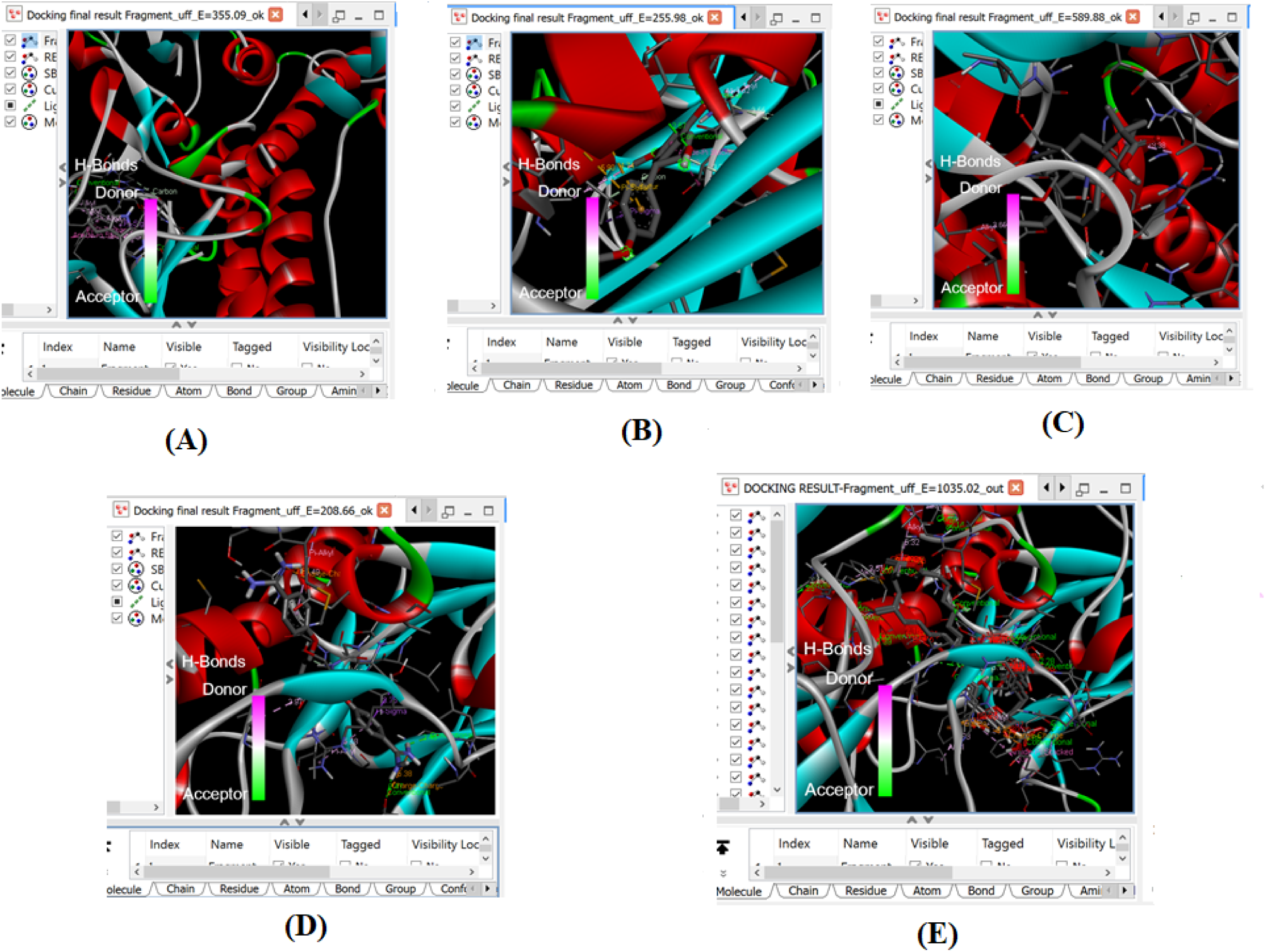
3D docking poses showing molecular interactions of harmane (A), abietic acid (B), epiafzelechin trimethyl ether (C), miltefosine (D) and amphotericin B (E) in the binding sites of trypanothione reductase.

## 3-2. DISCUSSION

Herbal plants are increasingly recognized as viable, sustainable, and cost-effective alternatives in the development of new anti-leishmanial agents, addressing the limitations of current treatments such as high toxicity, expense, and emerging drug resistance (Heidari-Kharaji et al., 2016). One such plant includes *Pinus* sp., which is used traditionally to treat fever conditions and other infections (Munteanu et al., 2025; Papp et al., 2022). This study evaluated the antileishmanial activity of extracts of *Pinus sp.* leaves. Since oxidative stress and inflammation are intricately involved in the pathogenesis of leishmaniasis, the antioxidant, anti-inflammatory and immunomodulatory effects of *Pinus sp.* extracts are also evaluated. Ethanol and methanol extracts from *Pinus sp.* leaves inhibited the growth of promastigotes (IC_50_ < 10 µg/mL) and amastigotes (IC_50_ < 25 µg/mL) of *L. donovani* with low IC_50_ values and high selectivity toward Raw (SI>50) cells. Previous reports have demonstrated that plant extracts with SI>10 are highly selective toward normal cells (Camacho et al., 2003; Indrayanto et al., 2021). Methanol, hydroethanol and hydromethanol extracts increased the NO production in infected macrophages, thus indicating the immunomodulatory effects of *Pinus sp.* extracts. As a matter of fact, lower concentrations of the plant extracts (˂20 µg/mL) decreased the levels of nitric oxide, whereas cell treatment with higher concentrations (100 µg/mL) increased the NO levels when compared to the untreated control cells. Moreover, *Pinus sp.* extracts revealed anti-inflammatory effects by inhibiting the denaturation of BSA with IC_50_ values ranging from 4.11 to 52.38 µg/mL, vs. diclofenac sodium (IC_50_: 2.91 µg/mL). Irrespective of the extract tested, *Pinus sp.* leaves showed significant antioxidant activity upon DPPH, ABTS and FRAP assays. The observed antileishmanial activity of *Pinus sp*. extracts might be attributed to their immunomodulatory, anti-inflammatory and antioxidant effects (Bhattacharya, 2025; Hassan et al., 2022; Mfopa Alvine Ngoutane & Kemzeu Raoul, 2024; Mfopa et al., 2024) Further phytochemical analysis of the most promising antileishmanial extract (Ps.FETOH) revealed the presence of phenolics and flavonoids. Accumulated evidence has shown the ability of plant derived flavonoids to suppress the release of pro-inflammatory cytokines by preventing protein misfolding and subsequent aggregation (Maspi et al., 2016; Rodrigues et al., 2025). Other researchers have demonstrated that plant-derived flavonoids prevent thermal denaturation of albumin, thus indicating anti-inflammatory effects (Enechi et al., 2020; Hasan et al., 2024; Precupas et al., 2016). The overproduction of nitric oxide following treatment with *Pinus sp*. extracts might have induced parasite cell apoptosis. Recently, it has been demonstrated that plant extracts can trigger high levels of NO, which directly damage the parasite machinery (Dhouafli et al., 2018). Previous studies indicate that certain flavonoids and phenolic compounds can induce parasite cell death by increasing the production of nitric oxide and triggering oxidative stress-mediated apoptosis (Mehwish et al., 2021; Sahakyan, 2026). While flavonoids are generally known as antioxidants, they can act as pro-oxidants in the presence of NO or specific metallic conditions, leading to the formation of reactive species that damage parasite DNA and mitochondrial function (Jomova et al., 2025).

Results obtained from the plant phytochemical screening are consistent with those of Singh and collaborators, who discovered high flavonoid and phenolic contents in *Pinus thunbergii* (Singh et al., 2020).

A single oral administration of the most promising extract (Ps.FEtOH) at 2000 and 5000 mg/kg in Wistar rats did not induce any signs of toxicity or mortality. After 14 days of observation period, and body weights, haematological (RBC, WBC, hematocrit, etc.) and biochemical parameters (ALT, AST, ALP, creatinine, etc.) of toxicity remained unchanged in treated and untreated groups of animals. The LD_50_ for the ethanol extract was found to be 5000 mg/kg. These results are consistent with the finding of Ghadirkhomi et al. (2016) who discovered a LD_50_ of 2000 mg/kg for the hydroethanolic extract of *Pinus eldarica*, a plant species from the genus *Pinus* (Ghadirkhomi et al., 2016).

The UHPLC-LCMS/MS analysis of Ps.FEtOH extract led to the identification of nine compounds, among which three compounds (abietic acid, epiafzelechin trimethyl ether and harmane) are well known for their occurrence in the *Pinus* genus (Fedorov & Ryazanova, 2021; Park et al., 2025; Roh et al., 2010; Zhang et al., 2025; Zhang et al., 2024). The observed antileishmanial activity of *Pinus* sp. extracts might be attributed to the presence of these secondary metabolites in the plant. Other compounds (alkylphenylketones, colensane and clerodane diterpenoids, and long-chain fatty acids) that were also identified are reported for their anti-inflammatory properties (Dewick, 2002; Duke, 2017; Hanson, 2011; Wilson et al., 2011). As already discussed, harmane (50 ≤ IC_50_ ≤ 100 µM; Di Giorgio et al., 2004) and epiafzelechin trimethyl ether (EC_50_: 13.6 µg/mL) inhibited the growth of *L. infantum* and *L. braziliensis*, respectively. Moreover, (Torres et al., 2020) reported the inhibitory effects of abietic acid towards selected *Leishmania* species (Torres et al., 2020). ADMET predictions revealed a favorable pharmacokinetic profile for harmane, abietic acid, and epiafzelechin trimethyl ether. Unlike the reference drug amphotericin B, these compounds did not violate any of Lipinski’s rules and revealed high oral bioavailability and intestinal absorption. However, harmane is predicted to have a high risk of toxicity. Further molecular docking studies predicted harmane (ΔG: −8.2 kcal/mol) as the compound with highest binding affinity to the target enzyme trypanothione reductase. At the active site of the complex harmane-trypanothione reductase, key active site residues ILE199 and TYR198 (hydrophobic) and SER162 and THR51 (H-bonds), suggesting a potent competitive inhibition (Bhardwaj et al., 2021; Kasahara et al., 2006; Saha et al., 2022). Although epiafzelechin trimethyl ether (ΔG:-7.8 Kcal/moL) and abietic acid (ΔG: −6.9 Kcal/moL) shows binding affinity close to that of miltefosine and amphotericin B, respectively, detailed mechanism of action of these compounds should be envisaged, because binding affinity do not dictate how the compound alters the target’s function or the kinetics of the molecular interaction (Decherchi & Cavalli, 2020; Tonge, 2018). The presence of the catalytic residues CYS52 and CYS57 at the binding pocket of the abietic acid-trypanothione reductase complex reveals their potential role to form a disulfide bridge (when oxidized) that is reduced to a dithiol during the catalytic cycle, allowing for the binding and reduction of trypanothione (Battista et al., 2020; Ilari et al., 2012).

## 4. LIMITATIONS AND PERSPECTIVES

This study summarizes the antileishmanial activity of *Pinus* sp. extracts against promastigote and amastigote forms of *L. donovani*. Extracts from ethanol, methanol, and from the mixtures water-ethanol (30:70; v/v) and water-methanol (30:70; v/v) inhibited the growth of *L. donovani* promastigotes (IC_50_s: 6.45-15.78 µg/mL) and amastigotes (IC_50_s: 16.02-24.44 µg/mL) with low IC_50_ values and high selectivity toward Raw cells (SI: >50), the most potent being the ethanol extract (**Ps.FEtOH**). Moreover, this extract demonstrates antioxidant and anti-inflammatory activities. Acute toxicity study revealed that **Ps.FEtOH** extract is nontoxic up to 5000 mg/kg, thus revealing a LD_50_>5000 mg/kg. The phytochemical screening of **Ps.FEtOH** revealed the presence of phenolic compounds and flavonoids. Further UHPLC-LCMS/MS analysis identified harmane, abietic acid and epiafzelechin trimethyl ether as the main compounds responsible for the antileishmanial effects. The prediction of ADMET properties of these compounds revealed their high bioavailability and intestinal absorption, even though harmane showed potential risks of toxicity. Further molecular docking of these compounds against trypanothione reductase as the target enzyme indicated highest binding affinity with harmane, followed by abietic acid and epiafzelechin trimethyl ether. This novel contribution on the inhibitory effects of *Pinus* sp. extracts toward *L. donovani* might help to identify potential leads for antileishmanial drug discovery. However, major limitations of this study include (i) isolation and characterization of antileishmanial compounds from *Pinus* sp. extracts; (ii) pharmacokinetic and *in vivo* tests, as well as detailed antileishmanial mechanisms of action.

Furthermore, extended studies on the inhibitory effects of extracts and compounds from *Pinus* sp. leaves toward other *Leishmania* species should be investigated to warrant the successful utilization of this plant in antileishmanial drug discovery.

## 5. CONCLUSION

This study demonstrates the antileishmanial effects of *Pinus sp.* extracts against promastigote and amastigote forms of *Leishmania donovani*. While non toxic upon acute oral toxicity testing (LD_50_>5000 mg/kg), the most active antileishmanial extract (Ps.FEtOH) revealed anti-inflammatory and immunomodulatory effects. The phytochemical analysis identified harmane, abietic acid, and epiafzelechin trimethyl ether as the potential compounds of *Pinus* sp. responsible for its biological action. Molecular docking of these plant compounds with trypanothione reductase revealed a high binding affinity of the ligands with the receptor. The prediction of the ADMET properties of the test compounds revealed favourable physicochemical and pharmacokinetic properties (bioavailability, intestinal absorption and solubility), even though harmane showed high risk of potential toxicity. Overall, the present study demonstrates the antileishmanial effects of *Pinus* sp. extracts. However, isolation and characterization of the active antileishmanial compounds from *Pinus* sp. extracts are warranted. Moreover, a detailed mechanism of action of antileishmanial extracts and compounds from *Pinus* sp., toxicity and *in vivo* experiments are needed to support the safe use of this plant in ethnomedicine.

## Acknowledgments

This work was carried out with funding from the YaBiNaPa (Yaounde-Bielefeld Graduate School for Natural Products with Antiparasite and Antibacterial Activity) project. We express our sincere gratitude to the project coordinator, Professor LENTA Ndjakou Bruno, for his valuable support and guidance throughout this study. We also thank Bei Resources for kindly providing the *Leishmania donovani* 1S strain (MHOM/SD/62/1S), NR-48821.

